# Notch signalling governs human enteric nervous system progenitor dynamics

**DOI:** 10.64898/2026.05.01.722150

**Authors:** Antigoni Gogolou, Nikolas Stefanidis, Fay Cooper, Guillaume Blin, Stanley E. Strawbridge, Alexander G. Fletcher, Anestis Tsakiridis

## Abstract

The enteric nervous system (ENS) is the main branch of the peripheral nervous system that innervates the gastrointestinal tract controlling vital functions. It arises during embryogenesis via migration, proliferation and differentiation of neural crest-derived ENS progenitors. Perturbation of these processes, caused by mutations in key signalling pathway components and transcription factors, prevents progenitor colonisation of the distal gut causing aganglionic phenotypes and enteric neuropathies such as Hirschsprung (HSCR) disease. While animal models implicate Notch signalling in ENS specification, its role in human ENS progenitor cell fate decisions remains unclear. Here, we employ a human pluripotent stem cell-based model to show that Notch signalling is a key regulator of human ENS progenitor dynamics. Quantitative modelling of our *in vitro* data indicates that Notch inhibition accelerates progenitor differentiation without substantially altering neuronal versus glial lineage bias. Furthermore, we establish that Notch signalling controls human ENS progenitor migration by influencing migration velocity and directionality. Together, these findings provide mechanistic insights into how Notch signalling disruption may contribute to the pathogenesis of human intestinal aganglionosis.

**SUMMARY STATEMENT:** *In vitro* generation of human enteric nervous system (ENS) cells and quantitative modelling reveal that Notch signalling regulates ENS progenitor differentiation rates and migration.

## INTRODUCTION

The gastrointestinal tract is primarily innervated by the intrinsic enteric nervous system (ENS), a complex network of neurons and glia organised into ganglia. The ENS controls several essential gut functions, including bowel motility, nutrient absorption and digestion, secretion of water, electrolytes, and hormones, and blood flow (Kang et al., 2021). It arises during embryonic development predominantly from the vagal neural crest (NC), a multipotent cell population specified at the neural plate border between somites 1 and 7 (Le Douarin and Teillet, 1973, Yntema and Hammond, 1954). After delaminating from the dorsal neural tube, vagal NC cells migrate into the foregut, while acquiring neuroglial-potent ENS progenitor features, and colonise the developing gut in a rostro-caudal direction.

The generation of appropriate proportions of enteric neurons and glia across space and time depends on the coordinated regulation of ENS progenitor migration, proliferation, and differentiation. Recent single-cell RNA-sequencing analysis in the mouse has challenged the long-held view that ENS progenitors remain uniformly bipotent throughout development, with equal propensity to generate enteric neurons and glia (Laddach et al., 2023). Instead, this study supports a model in which ENS progenitors are initially biased toward neurogenic differentiation and subsequently progress toward a default, quiescent gliogenic state. This branching of neurogenic differentiation trajectories from a linear gliogenic axis coincides with a progressive decline in progenitor proliferative capacity and migratory and colonisation potential (Zhang et al., 2019). Whether human ENS progenitors exhibit similar developmental dynamics remains unclear.

Disruption of ENS progenitor dynamics, caused by mutations in key signalling pathway components (e.g. *RET*, *EDNRB*) and transcription factors (e.g. *SOX10*), leads to failure of ENS progenitors to colonise the distal gut and, consequently, to aganglionic phenotypes and enteric neuropathies (Kang et al., 2021, Heuckeroth, 2018). Hirschsprung disease (HSCR) is the most representative of these conditions and is caused by the absence of enteric neurons in the distal bowel, which consequently loses propulsive motility, ultimately resulting in lethal intestinal obstruction (Heuckeroth, 2018). Thus, understanding how human ENS progenitor ontogeny is orchestrated at the molecular and cellular levels is critical for improving insight into the causes of enteric neuropathies.

One signalling pathway implicated in ENS development is Notch, a well-established and evolutionarily conserved modulator of cell fate. Notch receptors and their ligands, Delta and Jagged (in mammals), are transmembrane proteins that typically mediate interactions between adjacent cells. Ligand-receptor binding triggers a series of proteolytic events that ultimately lead to γ-secretase-dependent release of the Notch intracellular domain (NICD) from the cell membrane. NICD then translocates to the nucleus, where it cooperates with transcriptional effectors such as CBF1/RBP-Jκ to induce transcription of target genes, including the basic helix–loop–helix hairy and enhancer of split (HES) proteins (Bray, 2016).

In animal models, Notch signalling components are dynamically expressed in ENS progenitors, their derivatives, and the surrounding intestinal mesenchyme (Laddach et al., 2023, Sander et al., 2003, Liu and Ngan, 2014, Zhou et al., 2024, Jacobs-Li et al., 2023, Kuil et al., 2023, Morarach et al., 2021, McCallum et al., 2020). Moreover, deletion of Notch pathway members in mouse embryos disrupts multiple aspects of ENS development, including ENS progenitor maintenance, migration, proliferation, and differentiation (Mead and Yutzey, 2012, Okamura and Saga, 2008, Taylor et al., 2007, De Bellard et al., 2002). In line with these findings, expression of Notch pathway components is significantly perturbed in both animal and induced pluripotent stem cell (iPSC)-based models of HSCR, as well as in patient tissues (Vincent et al., 2023, Li et al., 2023, Ngan et al., 2011, Windster et al., 2025, Jia et al., 2012). Furthermore, genetic variant screening and genome-wide association studies have identified links between Notch-related mutations and HSCR (Ngan et al., 2011, Tang et al., 2017). However, the behaviour of human ENS progenitor cells under steady-state and disrupted Notch signalling conditions has not been comprehensively investigated.

We have previously established a protocol for the efficient generation of vagal NC cells exhibiting features of early ENS progenitors from human pluripotent stem cells (hPSCs) (Jevans et al., 2024, Gogolou et al., 2021, Frith et al., 2020). These cells were shown to efficiently colonise the ENS of immunocompromised mice, giving rise to neuronal and glial lineages following *in vivo* transplantation (Frith et al., 2020), and to improve contractility in aganglionic gut explants obtained from HSCR patients (Jevans et al., 2024). Here, we provide a detailed characterisation of the stepwise transition of these hPSC-derived vagal NC cells into later-stage ENS progenitors, enteric neurons, and glia *in vitro*, thereby establishing a robust model of human ENS development. Using this platform, we show that Notch signalling components are dynamically expressed during human ENS specification. Chemically inhibiting Notch signalling dramatically enhances the production of enteric neurons and glia. Computational modelling of our data, combined with compositional likelihood-based parameter inference and model selection, supports a branching differentiation hierarchy marked by an early pro-neurogenic bias. Our modelling further reveals that Notch inhibition alters developmental tempo, accelerating ENS progenitor differentiation and triggering the premature production of both neurons and glia. This effect is accompanied by increased neuronal clustering and reduced migratory capacity of ENS progenitors.

Together, our findings strongly support a critical role for Notch signalling in regulating the differentiation pace of human ENS progenitors. We propose that disruption in the temporal production of their neuronal and glial derivatives, coupled with impaired colonization potential, may underlie Notch-associated aganglionic phenotypes.

## RESULTS

### Characterisation of an *in vitro* model of human ENS development

We have previously defined culture conditions for the directed differentiation of hPSCs toward vagal NC cells and ENS cells (Frith et al., 2020, Gogolou et al., 2021). Our protocol involves the initial generation of an anterior NC progenitor population by day 4 of differentiation (D4) through combined stimulation of WNT signalling (via treatment with the GSK3 inhibitor CHIR99021), inhibition of TGFβ signalling, and moderate BMP activity. This is followed by the addition of retinoic acid (RA) to induce cells with vagal NC/early ENS progenitor features at D6 (**Fig. 1A**). D6 vagal NC/ENS progenitor cultures are then dissociated and cultured as non-adherent spheres in the presence of WNT, FGF, and RA signalling agonists for a further 3 days. The spheres are subsequently plated and cultured in neurogenic conditions to promote subsequent differentiation toward ENS components (**Fig. 1A**).

**Figure 1.**
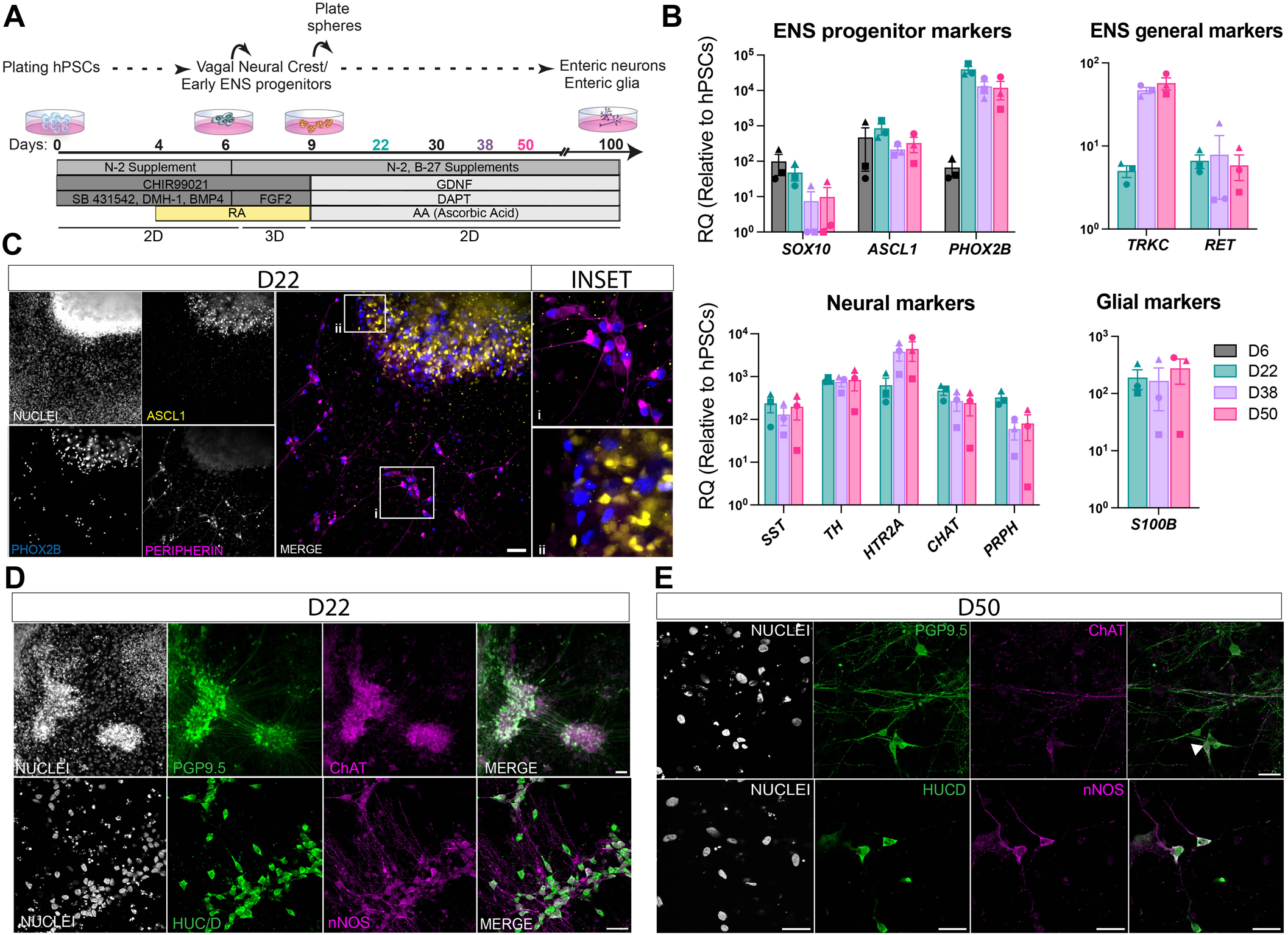
*In vitro* generation of enteric neurons from hPSCs. (A) Schematic of the protocol showing time points and culture conditions used to generate ENS cultures from hPSCs *in vitro*, including both two-dimensional (2D) and three-dimensional (3D) steps. (B) qPCR-based gene expression analysis of ENS progenitor, neuronal, and glial markers at the indicated time points. Data are shown as mean ± s.e.m., n=3 independent experiments with three replicates per experiment. D, differentiation day. (C) Immunofluorescence analysis of ASCL1, PHOX2B, and PERIPHERIN expression in D22 hPSC-derived ENS cultures. Insets show magnified regions of cells co-expressing PERIPHERIN and PHOX2B (i) and single ASCL1- and PHOX2B-positive cells (ii). (D, E) Immunofluorescence analysis of the indicated enteric neuron markers in D22 (D) and D50 (E) cultures. Scale bars: 50 μm (C), 30 μm (D, E).

We sought to characterize in detail the later stages of the protocol, after vagal NC specification, following sphere attachment and the differentiation of vagal NC-derived ENS progenitors into their derivatives. To this end, we differentiated hPSCs (the well-established iPSC line WTC11 (Miyaoka et al., 2014)) according to our protocol up to D50 (**Fig. 1A**). Time-course quantitative PCR (qPCR) analysis revealed that, following maximal induction at D6 (vagal NC stage), expression of the NC/ENS progenitor/glial marker *SOX10* progressively declined with differentiation time. In contrast, levels of the committed ENS progenitor/pro-neural regulators *ASCL1*, *PHOX2B,* and *RET* (Taraviras et al., 1999, Pattyn et al., 1999, Young et al., 2003, Memic et al., 2016) remained stable or increased between D22 and D50 (**Fig. 1B**). From D22 onwards, we detected induction of transcripts characteristic of enteric neurons (*TRKC, PRPH, TH*) (Chalazonitis et al., 2001, Morarach et al., 2021, Jacobs-Li et al., 2023, Szabolcs et al., 1996, Recinto et al., 2024), including markers of specific neuronal subtypes such as *SST* (interneurons) (Morarach et al., 2021), *HTR2A* (serotonergic neurons) (Schlieve et al., 2017), and *CHAT* (cholinergic neurons) (Sang and Young, 1998), as well as the glial marker *S100*β (Young et al., 2003, Ferri et al., 1982) (**Fig. 1B**).

We next interrogated the cellular composition of our cultures using immunofluorescence. At D10 (one day after sphere plating under ENS-inducing conditions; **Fig. 1A**) cells started migrating out of the spheres and differentiating into neurons, which subsequently formed clustered, interconnected neuronal networks (**Fig. S1A, S1B, 1C, 1D**), in line with other differentiation protocols employing a similar sphere formation step (Fattahi et al., 2016). By D22, PHOX2B protein was detected in the majority of PERIPHERIN (PRPH)^+^ neurons in low-density regions (**Fig. 1Ci**) and, in mutually exclusively with ASCL1, at the periphery of the re-plated spheres (**Fig. 1Cii**). In contrast, ASCL1^+^PHOX2B^-^ cells were largely confined to central regions within the spheres (**Fig. 1C**). These findings suggest a transition of ENS progenitors from an ASCL1⁺ to a PHOX2B⁺PRPH⁺ pro-neural state as they migrate out of the dense three-dimensional sphere environment.

The presence of nitrergic and cholinergic neurons at both D22 and D50 was confirmed by expression of neuronal nitric oxide synthase (nNOS) and choline acetyltransferase (ChAT), alongside pan-neuronal ENS markers HUC/D and PGP9.5 (Krammer et al., 1993, Desmet et al., 2014) (**Fig. 1D, E**). At D50, cultures contained PRPH⁺PHOX2B⁺ clusters, occasionally including ASCL1⁺ cells likely representing ENS progenitors committing to a neural fate (**Fig. S1C**). We also observed expression of the enteric neuron marker TRKC, while a subset of neurons exhibited immunoreactivity for tyrosine hydroxylase (TH), indicating a catecholaminergic character (**Fig. S1D, S1E**). A few cells expressed substance P, a neuropeptide found in subpopulations of excitatory neurons in the human ENS (Wattchow et al., 1988) (**Fig. S1F**). Finally, fibroblast-like cells were present in the cultures (as we recently reported in (Sabahi-Kaviani et al., 2026)), potentially reflecting the mesenchymal potential of the vagal NC/ENS progenitors (Ling and Sauka-Spengler, 2019, Bixby et al., 2002) and/or possible contamination by cranial NC derivatives. We also investigated the induction of ENS progenitors and glial cells in our cultures. In the mouse embryo, early ENS progenitors co-express SOX10 and CD49d (ITGA4), which also mark their default glial character acquired during later ENS development (Lasrado et al., 2017, Laddach et al., 2023). This transition toward a gliogenic state is accompanied by the upregulation of more definitive glial markers such as S100β. Mapping of SOX10^+^ and CD49d^+^ cells using immunofluorescence analysis revealed that double-positive cells were already present at D11 (two days after sphere plating) and persisted throughout the differentiation period examined up to D22 (**Fig. 2A**). Neuronal commitment, reflected by PGP9.5 expression, was apparent at D11 and coincided with the loss of CD49d expression, indicating exit from the progenitor state (**Fig. 2B**). At later differentiation stages (D22 and D50), SOX10^+^CD49d^+^ cells upregulated S100β expression, consistent with a transition to a glial fate; these glial cells were interspersed among TUJ1⁺ neurons (**Fig. 2C-E, S2C**). Similar results were obtained using a second independent hPSC line, a WA09/H9 human embryonic stem cell (hESC)-derived SOX10:GFP line (Chambers et al., 2012) (**Fig. S2**).

**Figure 2.**
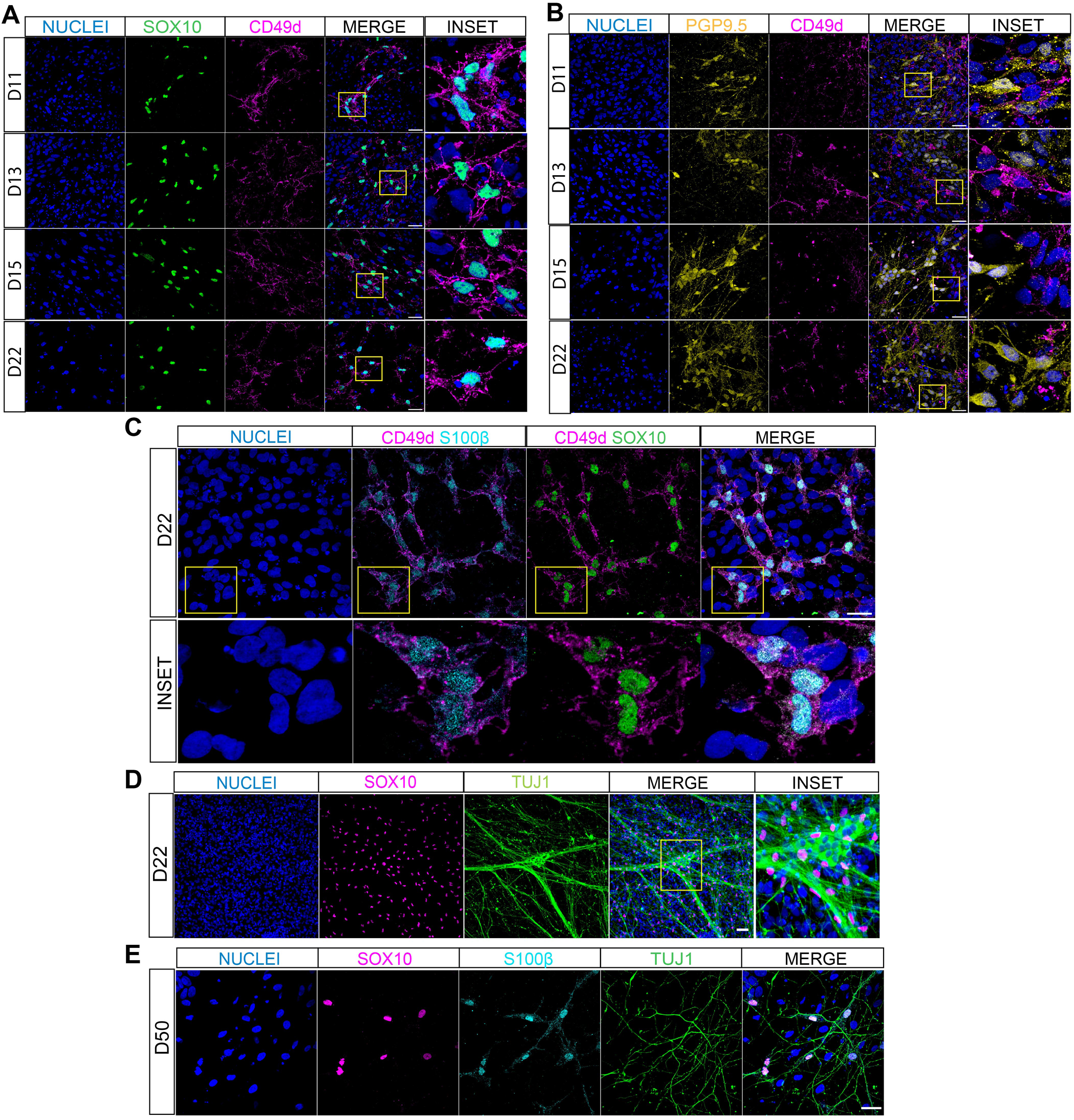
*In vitro* generation of ENS progenitors and glia from hPSCs. (A, B) Immunofluorescence analysis of the expression of CD49d together with the ENS progenitor/glial marker SOX10 (A) or the neural marker PGP9.5 (B) at the indicated differentiation time points (D, days). Insets show magnified areas. (C) Immunofluorescence analysis of the expression of CD49d together with the glial markers SOX10 and S100β in D22 cultures. Magnified regions of the indicated areas are shown in the lower panel. (D, E) Immunofluorescence analysis of the expression of neuronal (TUJ1) and glial (SOX10, S100β) markers in D22 (D) and D50 (E) cultures. Scale bars: 30 μm (A-C, E) and 50 μm (D).

Together, these results demonstrate that our differentiation protocol successfully produces a diverse population of enteric neurons and glial cells from hPSCs.

### Dynamics of Notch signalling component expression during the induction of human ENS cells

The final sphere-plating step of our protocol includes the addition of the Notch signalling/γ-secretase inhibitor DAPT (**Fig. 1A**), based on our observation that its inclusion significantly enhances the yield of ENS cells derived from hPSCs and consistent with previous reports on other neural differentiation protocols (Frith et al., 2020, Maury et al., 2015, Borghese et al., 2010). Given the established role of Notch signalling in mouse and zebrafish ENS development and its association with HSCR pathogenesis, we examined the effects of Notch inhibition in our human ENS differentiation model, comparing DAPT-treated cultures to DMSO-treated controls (**Fig. 3A**). To minimise potential off-target effects, we used the lowest effective concentration of DAPT (10 µM), which was sufficient to reduce the expression of canonical Notch target genes in hPSC-derived ENS cultures (**Fig. S3A, B**).

**Figure 3.**
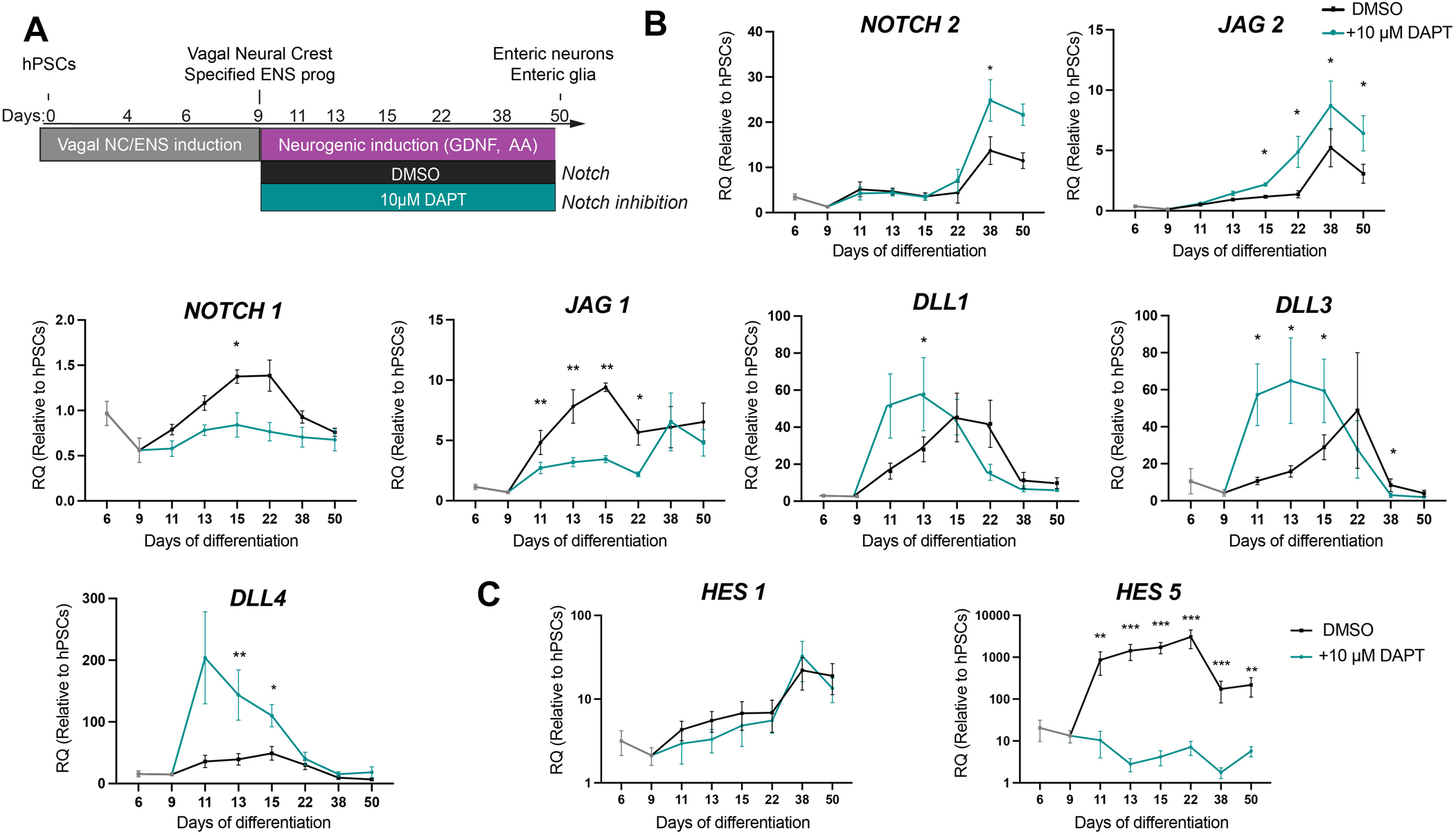
Dynamics of Notch signalling component expression during human ENS differentiation. (A) Schematic of the time points and culture conditions used to generate ENS cultures with or without Notch pathway inhibition. (Β) Time-course qPCR analysis of the indicated Notch signalling components in DMSO control (black line) and DAPT-treated (green line) ENS cultures. Data are presented as mean ± s.e.m., n=5 independent experiments with three replicates per experiment; *P <0.05, **P <0.01, ***P<0.001 (paired t-test). Only statistically significant differences are indicated.

We first assayed the dynamics of Notch signalling component expression in our differentiating ENS cultures from D6 to D50 under control (DMSO) conditions using qPCR. We observed two distinct expression patterns: high *NOTCH1*, *JAG1*, *DLL1*, *DLL3* and *DLL4* transcript levels were prevalent during the early phase of ENS differentiation (D13–22), suggesting a role in ENS progenitor cell fate decisions and early lineage commitment; in contrast, *NOTCH2* and *JAG2* expression increased during the later differentiation phase (D22–50), suggesting a role in committed enteric neurons and glia (DMSO control, black line, **Fig. 3B**). These findings are consistent with previous reports showing enrichment of *NOTCH1*, *DLL1*, and *DLL3* transcripts in mouse ENS progenitors (Laddach et al., 2023, Morarach et al., 2021, Okamura and Saga, 2008). We also analysed the Notch target genes *HES1* and *HES5*. *HES5* expression peaked during the early progenitor-associated period, whereas *HES1* reached maximal levels at later differentiation stages (black line, **Fig. 3C**). DAPT treatment markedly reduced *HES5* expression, confirming effective Notch pathway inhibition, while *HES1* levels were largely unaffected (**Fig. 3C**, green vs black lines). Furthermore, DAPT treatment significantly downregulated *NOTCH1* and *JAG1* (**Fig. 3B**), while triggering upregulation in *DLL1/3/4*, which also exhibited earlier expression peaks compared with DMSO controls (**Fig. 3B**).

Collectively, these findings suggest that Notch signalling is active during hPSC-derived ENS cell specification and is regulated by the dynamic expression of distinct receptor and ligand-encoding genes, which exhibit differential temporal sensitivity to DAPT-mediated inhibition.

### Notch signalling inhibition enhances human ENS progenitor differentiation

We next sought to define the impact of DAPT-mediated Notch signalling perturbation on ENS cell induction. Time-course qPCR analysis showed that DAPT treatment resulted in upregulation of both pro-neural (*ASCL1, PHOX2B*, *PRPH*) and glial (*SOX10, S100*β)-associated transcripts at different time points throughout the course of differentiation (**Fig. 4A, B**). This was accompanied by an increase in the expression of enteric neuron subtype markers such as *SST, TH*, and *CHAT,* by D22 (**Fig. S3C**).

**Figure 4.**
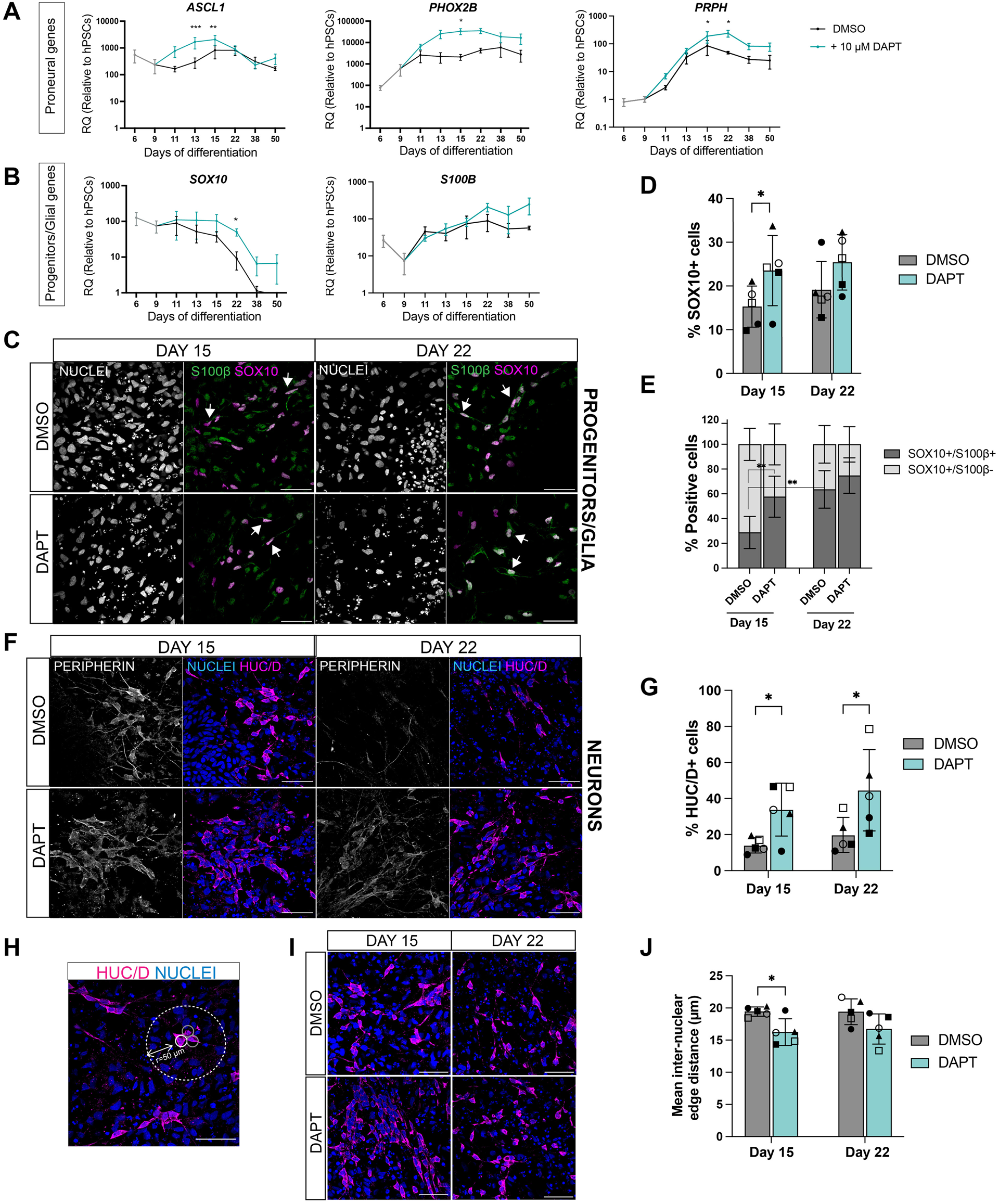
Notch signalling inhibition enhances human ENS progenitor differentiation. (A, B) Time-course qPCR analysis of the indicated proneural (A) and ENS progenitor/glial (B) genes in DMSO control (black line) and DAPT-treated (green line) cultures. Data are shown as mean ± s.e.m, n=5 independent experiments with three replicates per experiment. *P <0.05, **P <0.01, *** P<0.001 (paired t-test). (C) Immunofluorescence analysis of SOX10 and S100β expression in control and DAPT-treated ENS cultures at days 15 and 22. (D) Quantification of the percentage of SOX10^+^ cells per field of view at days 15 and 22. Data are presented as mean ± s.d., n = 5 independent experiments, with 10-15 fields scored per condition, time point, and experiment. Each biological replicate is represented by a unique symbol. *P <0.05 (paired t-test). (E) Quantification of double-positive (SOX10^+^S100 β^+^; glia) and single-positive (SOX10^+^S100^-^; ENS progenitors) cells in control and DAPT-treated cultures at days 15 and 22. Stacked bars represent mean ± s.d., n=5 independent experiments, with 15-30 fields scored per condition, time point, and experiment; **P <0.01, (paired t-test for the indicated comparisons). (F) Immunofluorescence analysis of PERIPHERIN and HUC/D expression in control and DAPT-treated ENS cultures at days 15 and 22. (G) Quantification of the percentage of HUC/D^+^ cells per field of view at days 15 and 22. Data are presented as mean ± s.d., n = 5 independent experiments, with 30 fields scored per condition and experiment for day 15 and 15 fields scored per condition and experiment for day 22. Each independent experiment is represented by a unique symbol. *P <0.05 (paired t-test). (H) Method for measuring internuclear edge distances. Analysis was performed on all HUC/D^+^ cells. As cytoplasmic boundaries were limited, nuclei were used as proxies to identify nearest neighbours. Using Delaunay triangulation, each nucleus (white outline) was paired with neighbouring nuclei (grey outline) within a 50 µm radius (white dotted line), and internuclear edge distances were calculated across the image. (I, J) Representative immunofluorescence images of HUC/D^+^ cells (I) and quantification of mean internuclear edge distances for at days 15 and 22 (J). Data are shown as mean ± s.d., n=5 independent experiments, each containing hundreds to thousands of HUC/D^+^ cells. Each biological replicate is represented by a unique symbol; *P <0.05 (paired t-test). Only statistically significant comparisons are indicated. Arrows indicate double-positive cells. Scale bars: 50 μm.

To further examine the effects of DAPT treatment on ENS progenitor dynamics, we assessed proteins marking ENS progenitor (SOX10^+^S100β^-^), neuronal (HUC/D, PERIPHERIN), and glial (SOX10^+^S100β^+^) identities at early (D15) and late (D22) differentiation stages. Under DMSO control conditions, the proportion of SOX10^+^S100β^+^ glial cells increased markedly from D15 to D22 (mean 64±15% in D22 vs 29±13% in D15, P<0.01) (**Fig. 4C-E**; dark grey bars in **4E**), possibly reflecting the default gliogenic fate of ENS progenitors, consistent with mouse studies (Laddach et al., 2023). In contrast, the production of HUC/D+ neurons remained relatively constant between these same two timepoints (**Fig. 4F, G**; dark grey bars in **4G**).

Consistent with our previous observations (Frith et al., 2020), DAPT treatment significantly increased the percentage of HUC/D^+^ cells compared with DMSO controls at both D15 (mean 34±15% in DAPT-treated vs 14±4% in control; P<0.05) and D22 (mean 45±23% in DAPT-treated vs 20±10% in control; P<0.05) (**Fig. 4F, G**). Additionally, DAPT treatment significantly increased the percentage of SOX10^+^ cells at D15 and, to a lesser degree, at D22 (**Fig 4C, D**). This expanded population consisted predominantly of SOX10^+^S100β^+^ glial cells (58±17% in DAPT vs 29±13% in control, P<0.01), with a corresponding reduction in SOX10⁺S100β⁻ ENS progenitors (**Fig. 4C, E)**. As all S100β⁺ enteric glial cells co-express SOX10, S100β single-positive cells, likely representing non-glial contaminants, were excluded from analysis (Hernandez-Ortega et al., 2024, Boesmans et al., 2015)). The effect of DAPT on progenitor differentiation dynamics was also accompanied by enhanced neuronal clustering at D15, as revealed by nearest neighbour analysis showing a significantly reduced mean internuclear edge distance in DAPT-treated samples (**Fig. 4H-J**).

Collectively, these data demonstrate that disruption of Notch signalling via the γ-secretase inhibitor DAPT enhances human ENS progenitor differentiation toward both neuronal and glial lineages.

### Quantitative modelling supports a role for Notch signalling in regulating the pace of ENS progenitor differentiation

Extensive evidence from various developmental systems has indicated that Notch signalling is a key regulator of differentiation timing (Gonzalez and Reinberg, 2025, Bray and Bigas, 2025). We therefore employed a compartment modelling framework to evaluate how ENS progenitor differentiation rates and lineage biases evolve in our *in vitro* model, both with and without Notch inhibition (**Fig. 5A-C**). Using a compositional likelihood-based approach, we inferred parameter sets for which our simulations sufficiently match the empirical data (**Fig. 4E, G**). We tested four distinct scenarios: constant differentiation rate and lineage biases (CC); constant rate and time-varying biases (CV); time-varying rate and constant biases (VC); and time-varying rate and biases (VV). This approach aligns with (Laddach et al., 2023), which identified a temporal neurogenic-to-gliogenic shift in the lineage bias of mouse ENS progenitors.

**Figure 5.**
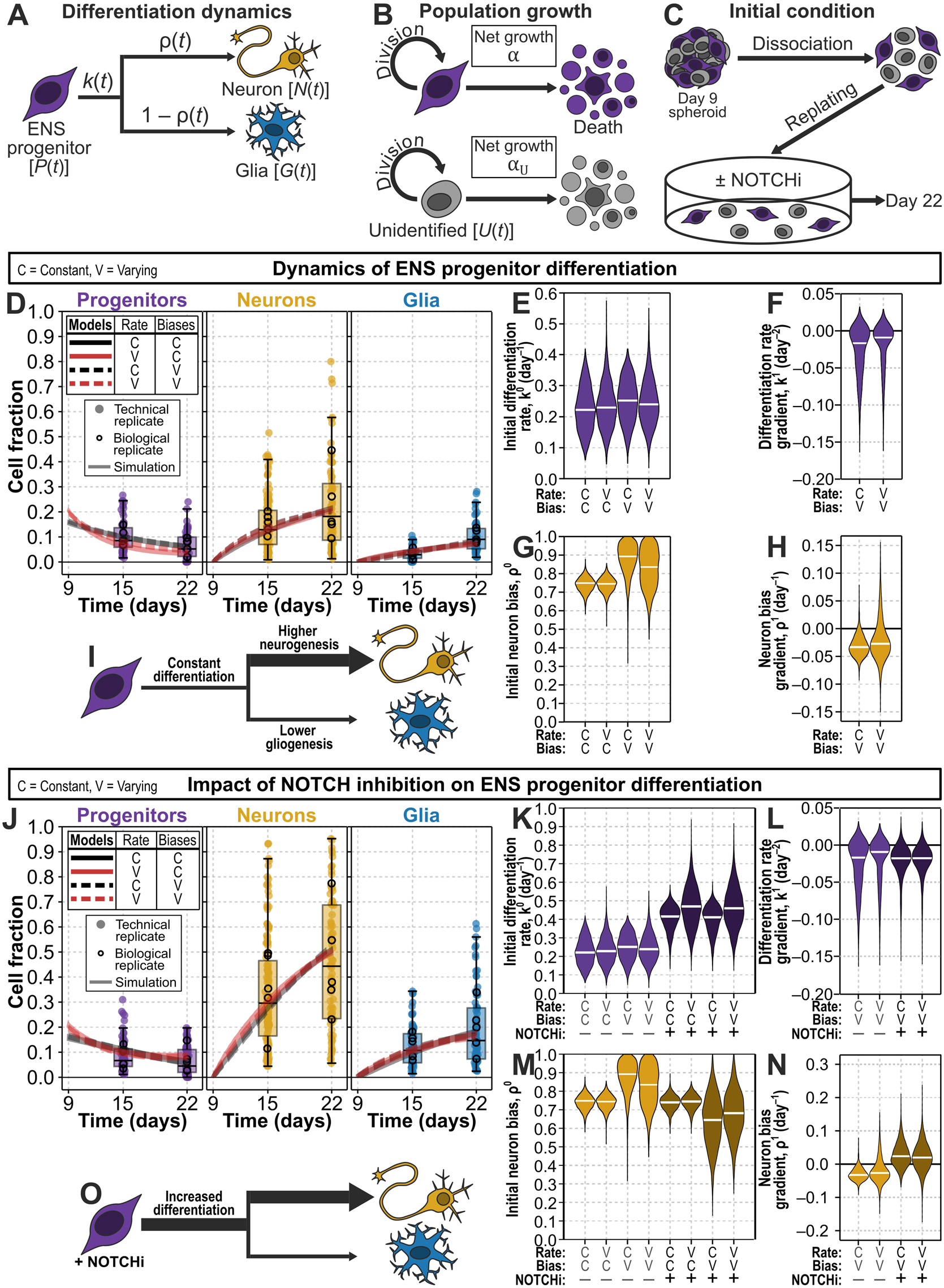
Quantitative modelling supports a role for Notch signalling in regulating the pace of human ENS progenitor differentiation. (A) Our modelling of ENS progenitor differentiation *in vitro* assumes that ENS progenitors, *P* (SOX10^+^S100β^-^) differentiate at per-capita rate *k* into neurons, *N* (HUC/D,), or glial cells, *G* (SOX10^+^S100β^+^), with neuronal and glial shares *ρ* and 1- *ρ*, respectively. An unidentified population, *U* (SOX10⁻S100β⁻HUC/D⁻), co-emerges in the cultures but is not derived from the ENS progenitors. (B) ENS progenitors proliferate with net per-capita growth rate *α*, which takes into account both cell division and cell death; glia are produced by differentiation and do not appreciably self-renew; the unidentified population evolves with its own net per-capita rate *α_U_*; and neurons are quiescent. (C) Initial conditions following dissociation and replating at day 9 of the differentiation protocol. (D) Numerical solutions using median posterior parameters from all four models (lines) fitted to empirical data on ENS progenitor differentiation dynamics (markers and boxplots; see also **Fig 4E, G,** DMSO). Filled circles indicate technical replicates, open circles indicate biological replicates, and box plots show the median and interquartile range of technical replicate data. (E-H) Posterior distributions of initial differentiation rates (E), differentiation rate gradients (F), initial lineage biases (G), and lineage bias gradients (H) for all four models. (I) Summary schematic of inferred ENS progenitor differentiation dynamics. (J) Numerical solutions using median posterior parameters from all four models (lines) re-fitted to empirical data on ENS progenitor differentiation dynamics under NOTCH inhibition (markers and boxplots; see also **Fig 4E, G**, DAPT). Filled circles indicate technical replicates, open circles indicate biological replicates, and box plots show the median and interquartile range of technical replicate data. (K–N) Posterior distributions of initial differentiation rate (K), differentiation rate gradient (L), initial lineage bias (M), and neuronal lineage bias gradient (N) when re-fitted to NOTCH inhibition data. (O) Summary schematic for the inferred impact of NOTCH inhibition on ENS progenitor differentiation dynamics.

We first examined normal ENS progenitor differentiation using our SOX10, S100β and HUC/D expression data from D15 and D22 cultures. (**Fig. 4E, G**; DMSO). Simulated dynamics for the median posterior parameter values showed high fidelity to the observed data (**Fig. 5D, S4A-D**). Leave-one-biological-replicate-out cross-validation did not identify a single preferred scenario, with differences in expected log predictive density between models small relative to their uncertainty (all within approximately two standard errors of the constant-rate, constant-bias model). This suggests that current experimental resolution is insufficient to definitively distinguish between these scenarios.

Analysis of inferred median posterior parameters across all models (**Fig. S4E**) showed comparable differentiation rates across scenarios (CC: 0.221 [0.102, 0.368]; VC: 0.226 [0.111, 0.397]; CV: 0.250 [0.113, 0.374]; VV: 0.242 [0.118, 0.403] day⁻¹; median and 95% credible interval), with all pairwise credible intervals overlapping across ≥94% of the narrower interval (**Fig. 5E**). In the time-varying models the rate gradient was weakly identified, with credible intervals spanning zero (VC: −0.016 [−0.107, +0.016]; VV: −0.009 [−0.097, +0.015] day⁻²) and posterior contraction relative to the prior of only 55% and 64% respectively (**Fig. 5F**). At D22, the inferred differentiation rate was 0.000 day^-1^ [0.000, 0.424] (VC) and 0.103 day⁻¹ [0.000, 0.441] (VV), computed per posterior draw as *k*^0^ + 13*k*^1^ and floored at zero as in the forward model; the constant-rate models give 0.221 [0.102, 0.368] (CC) and 0.250 [0.113, 0.374] (CV) day⁻¹ by construction. The breadth of these intervals reflects the weak identification of the gradient rather than evidence for a low terminal rate. In summary, the data do not resolve a temporal trend in differentiation rate over this interval. We therefore adopted the constant-rate models for subsequent analysis on grounds of parsimony, noting that a declining rate is not excluded.

Regarding lineage biases, all models inferred a strong and consistent preference for the neuronal over the glial lineage (neuronal share of differentiation CC: 0.741 [0.658, 0.811]; VC: 0.741 [0.660, 0.811]; CV: 0.894 [0.683, 0.993]; VV: 0.837 [0.608, 0.990]; median and 95% credible interval; **Fig. 5G**), reflecting previous reports of early neuronally-biased ENS progenitors (Laddach et al., 2023). In the time-varying models this neuronal share declined over time (CV: −0.033 [−0.063, +0.008] day⁻¹, posterior probability of a decline 0.95; VV: −0.029 [−0.084, +0.074] day⁻¹, probability 0.80; **Fig. 5H**), reaching 0.463 [0.141, 0.818] (CV) and 0.485 [0.000, 1.000] (VV) by D22, computed per posterior draw as ρL + 13ρ¹. In the CV model the decline is comparatively well resolved (86% posterior contraction relative to the prior) and carries the neuronal share across the 0.5 threshold, providing evidence for a time-dependent shift from neuronal towards glial production, consistent with previous work (Laddach et al., 2023); in the VV model, where the rate gradient is estimated simultaneously, it is not resolved. Within this timeframe our data are nonetheless equally well described by a constant differentiation rate and a neuronally-biased fate choice (**Fig. 5I**), which we adopted for subsequent analysis.

We next applied this framework to quantify kinetic shifts under Notch inhibition. As with the control, simulated dynamics (**Fig. 5J**, **S5A-D**) showed high fidelity to the data (**Fig. 4E, G**; DAPT), and again no single scenario was preferentially supported by cross-validation. Consistent trends emerged across the median posterior parameters (**Fig. S5E**). Notch inhibition increased the initial differentiation rate in all four models (CC: 0.415 [0.248, 0.514]; VC: 0.468 [0.257, 0.703]; CV: 0.408 [0.234, 0.511]; VV: 0.463 [0.266, 0.703] day⁻¹; median and 95% credible interval; **Fig. 5K**), with posterior probability of an increase relative to control exceeding 0.92 in every model and median estimates corresponding to an approximately two-fold change (pooled ratio 1.84 [0.88, 4.27]), though the magnitude of the increase is imprecisely determined. In the time-varying models, the rate gradient was of similar magnitude to control (VC: −0.018 [−0.079, +0.010]; VV: −0.019 [−0.082, +0.010] day⁻²; **Fig. 5L**), with credible intervals spanning zero and no detectable difference between conditions (VC: +0.001 [−0.067, +0.092]; VV: −0.007 [−0.072, +0.081] day⁻²). Notch inhibition therefore elevates the differentiation rate without detectably altering its temporal trend over this interval.

In contrast to the differentiation rate, the neuronal vs glial bias was essentially unchanged under Notch inhibition: in the bias-constant models the neuronal share of differentiation remained close to its control value (CC: 0.739 [0.648, 0.809] vs 0.741; VC: 0.743 [0.657, 0.810] vs 0.741; posterior probability of any between-condition difference 0.49 and 0.51 respectively; **Fig. 5M**), indicating that Notch inhibition accelerates differentiation without substantially altering the initial balance between neuronal and glial production. In the bias-varying models, however, the temporal trend differed in sign from control, the neuronal share increasing rather than declining over the interval (CV: +0.024 [−0.034, +0.105]; VV: +0.020 [−0.052, +0.106] day⁻¹; **Fig. 5N**), reaching 0.948 [0.429, 1.000] (CV) and 0.935 [0.213, 1.000] (VV) by D22. The between-condition difference in this gradient was suggestive but not decisive (CV: +0.057 [−0.014, +0.143] day⁻¹, posterior probability 0.94; VV: +0.047 [−0.073, +0.150] day⁻¹, probability 0.82), and we therefore interpret it cautiously: the data are consistent with Notch inhibition sustaining, rather than progressively reducing, the neurogenic bias over this window. The pronounced between-condition shift in lineage bias reported previously was not supported once the unidentified cells were treated as a co-emerging population rather than a progenitor-derived fate.

Because the SOX10^−^/S100β^−^/HUC/D^−^ co-emerging population is dynamically uncoupled from the ENS lineages in our model, the composition of the ENS compartment is independent of both α_U and P₀, and the parameter estimates and model comparison above are unaffected by how this population is treated. We therefore also present the data of **Fig. 4E, G** normalised to the ENS compartment alone (SOX10⁺ progenitors, HUC/D⁺ neurons and S100β⁺ glia; **Fig. S6**). This presentation removes the effect of the declining unidentified fraction, which differs markedly between conditions (model-inferred median falling from 0.85 to 0.65 between D9 and D22 in control, and from 0.84 to 0.27 under Notch inhibition). Normalised in this way, the apparent increase in neuronal proportion in control between D15 and D22 (1.44-fold when expressed as a fraction of all cells) is largely attributable to this dilution effect, with the neuronal share of the ENS compartment essentially unchanged (0.54 to 0.53); the principal change within the compartment is a shift from progenitors to glia (0.35 to 0.20 and 0.12 to 0.27 respectively), consistent with the inferred lineage-bias dynamics. Corresponding values under Notch inhibition were 0.60 to 0.63 (neurons), 0.15 to 0.10 (progenitors) and 0.24 to 0.27 (glia).

This decline in the unidentified population is captured directly by its inferred net per-capita rate, which was indistinguishable from zero in control (*α_v_* = −0.019 [−0.133, +0.079] day⁻¹) but negative under Notch inhibition (−0.121 [−0.241, −0.043] day⁻¹; posterior probability of a decline 1.00 in all four models), the between-condition difference being −0.103 [−0.254, +0.037] day⁻¹ (posterior probability 0.92; **Fig. S4E, S5E**). This population was the only one whose proportion decreased following Notch inhibition: comparing conditions at D22, the proportions of neurons and glia among all cells scored were higher under Notch inhibition (0.223 vs 0.456 and 0.102 vs 0.196 respectively; empirical biological-replicate means) and that of progenitors essentially unchanged (0.066 vs 0.067), whereas the progenitor share of the ENS compartment alone fell (0.195 vs 0.097; **Fig. S6**), consistent with accelerated differentiation.

Overall, Notch inhibition accelerates ENS progenitor differentiation increasing the production of both neurons and glia while leaving the neuronal vs glial bias largely stable, consistent with the branching model proposed recently (Laddach et al., 2023) in which an early neurogenic phase precedes a later gliogenic phase (**Fig. 5O**). These findings indicate that Notch signalling is a key regulator of the differentiation pace in ENS progenitors, ensuring the temporally refined production of neuronal and glial derivatives.

### Notch-inhibited ENS progenitors exhibit impaired migration capacity

As ENS progenitor differentiation and migration are tightly coupled processes (Zhou et al., 2024, Uesaka et al., 2008, Baker et al., 2022, Young et al., 2004, Young et al., 2014, McKeown et al., 2017), and considering that Notch inhibition induces both accelerated differentiation (**Fig. 5**) and HUC/D⁺ neuron clustering at D15 (**Fig. 4**), we next investigated whether Notch signalling disruption also influences migratory behaviour. To this end, ENS cultures were treated with either DAPT or DMSO at the sphere plating stage (D9), followed by a scratch assay at D14. Wound closure was then monitored over a 24-hour period (up to D15) using live imaging (**Fig. 6A**).

**Figure 6.**
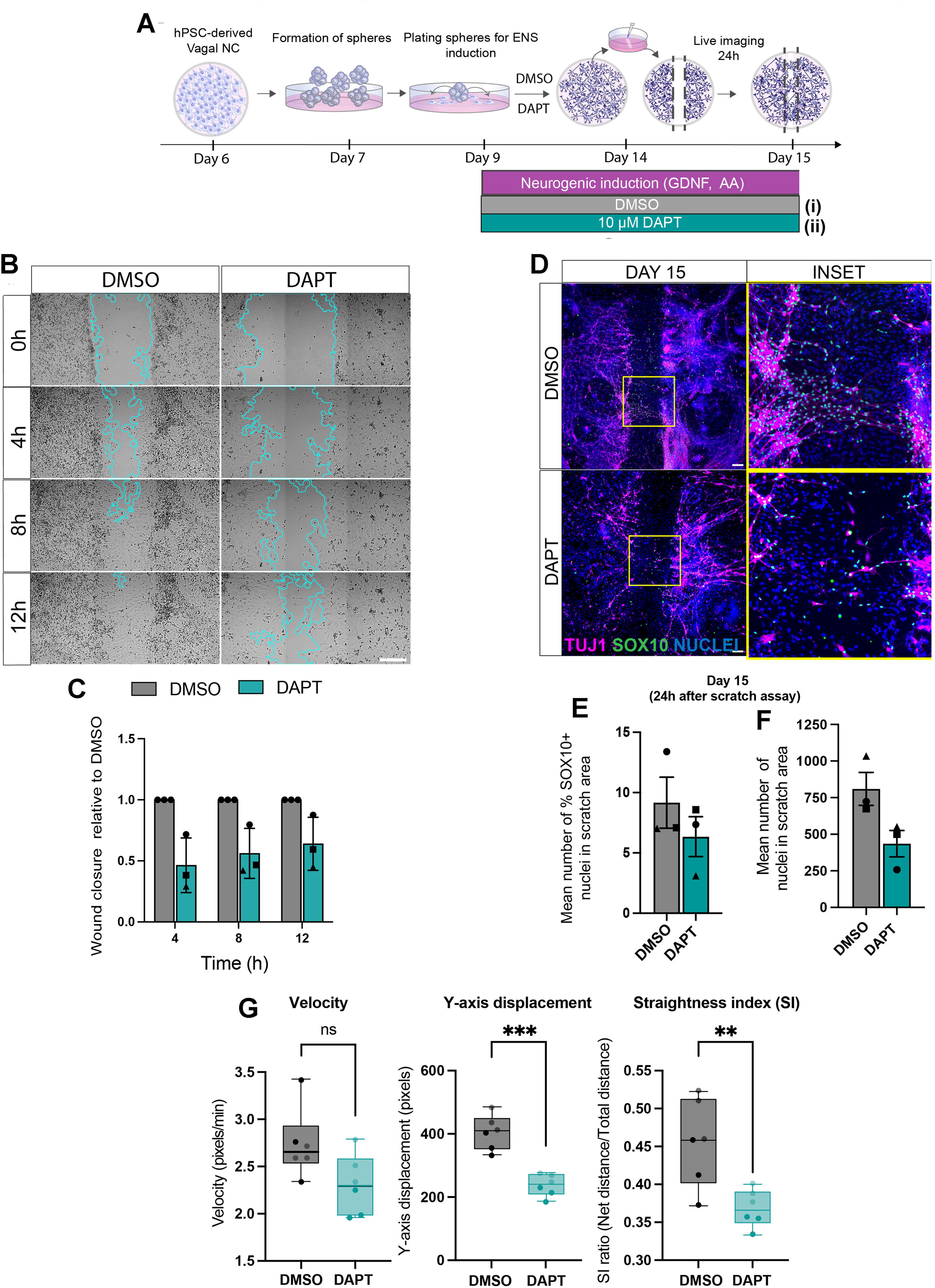
Notch signalling inhibition impairs human ENS progenitor migration. (A) Schematic of the scratch assay. hPSC-derived vagal NC cells formed into spheres (day 6) and plated for ENS induction (day 9). By day 14, cells formed a confluent monolayer, after which a scratch was introduced using a pipette tip. Cell migration into the wound was monitored by live imaging. Notch pathway conditions were: (i) DMSO control added at day 9 (grey bars); (ii) DAPT added at day 9 (green bars). (B) Time-lapse images of a representative scratch assay for DMSO- and DAPT-treated ENS cultures at 0, 4, 8, and 12 hours post-scratch. Leading edges were detected using a segmentation algorithm (blue lines). Scale bar: 200 μm. (C) Wound closure relative to DMSO control (set as 1). Data are shown as mean ± s.d., n=3 independent experiments with 3-5 fields scored per condition. Fields were sampled from 1-3 wells per experiment. *P <0.05 (unpaired t-test). (D) Immunofluorescence analysis of SOX10 and TUJ1 in DMSO- and DAPT-treated ENS cultures at day 15 (24 hours post-scratch). Insets show magnified regions. Scale bar: 200 μm. (E, F) Mean percentage of SOX10^+^ cells and mean number of total nuclei in the wound area 24 hours post-scratch. Data are shown as mean ± s.e.m, n=3 independent experiments with 4-10 fields scored per experiment. Each biological replicate is represented by a unique symbol. (G) Quantification of ENS cell migration parameters in the presence of DAPT/DMSO following live imaging of wound closure (see **Fig. S7C**): cell migration velocity (pixels/min) (left), Y-axis displacement (pixels) representing directional migration toward the wound centre over time (centre), and Straightness index (ratio of net displacement to total path length) measuring directional persistence during wound healing (right). Data are shown as box-and-whisker plots displaying median, interquartile range and minimum-to-maximum values. Data points represent n=2 biological repeats per condition (light and dark points), 3 fields scored per condition, 10 cells per field. *** P< 0.001, ** P< 0.01 (unpaired t-test).

DAPT treatment resulted in a delay in wound closure compared with DMSO-treated control ENS cultures (**Fig. 6B, C, S7A,B**). Immunostaining of scratch assay cultures at D15 revealed that wound closure in DMSO control cultures involved chain-like migration of SOX10^+^ progenitors within the wound space and their association with nascent TUJ1^low/+^ neuronal networks bridging the opposing wound sides (**Fig. 6D**). This may reflect the previously reported upregulation of neuronal markers in ENS progenitors during transition toward neural commitment (Zhou et al., 2024, Hao et al., 2009, Bondurand et al., 2006, Barlow et al., 2003). In contrast, DAPT treatment severely disrupted this process, evidenced by the absence of neuronal bridges and a reduction in the total number of cells and percentage of SOX10⁺ ENS progenitors within the wound region (**Fig. 6D-F**). Further analysis of live imaging data quantifying single-cell migration dynamics during wound closure showed that DAPT-treated ENS cultures exhibited reduced migration velocity and, most prominently, migration directionality compared with DMSO-treated controls (**Fig. 6G, S7C**).

Together, these findings indicate that the intrinsic migratory behaviour of early ENS progenitors is influenced by Notch signalling and is likely linked to the timing of their differentiation toward neurons and glia.

## DISCUSSION

Several studies using animal models have revealed a multifaceted role for Notch signalling in controlling both ENS and NC development, encompassing progenitor maintenance, proliferation, migration, and differentiation (Mead and Yutzey, 2012, Taylor et al., 2007, Okamura and Saga, 2008, Wakamatsu et al., 2000, Morrison et al., 2000, ME et al., 2007, Delfino-Machin et al., 2017, Wiszniak and Schwarz, 2019, Okubo et al., 2021, Alhashem et al., 2022). These functions can be opposing, depending on developmental stage and specific ligand-receptor interactions. The importance of Notch is further highlighted by evidence that the expression of many of its components is perturbed in both animal models and patients with HSCR, a potentially fatal enteric neuropathy resulting from lack of innervation of the distal bowel (Vincent et al., 2023, Li et al., 2023, Ngan et al., 2011, Windster et al., 2025, Tang et al., 2017, Jia et al., 2012). Despite this, the role of Notch signalling in human ENS progenitor ontogeny remains largely unexplored.

To address this, here we characterised an *in vitro* model of human ENS development involving the stepwise differentiation of hPSCs into ENS progenitors, which subsequently give rise to enteric neurons (including multiple subtypes) and glia (**Fig. 1, 2**). To interrogate how the proportions of emerging enteric neurons and glia shift over time, we coupled our platform with computational modelling. Our results align with previous single-cell RNA-sequencing studies in mice, which suggest that early ENS progenitors are biased toward neurogenesis (**Fig. 5**) (Laddach et al., 2023). Crucially, this approach revealed that Notch signalling regulates the pace of ENS progenitor differentiation: DAPT-mediated inhibition led to a marked increase in both neuronal and glial numbers at the expense of the progenitor pool, driven by an accelerated rate of differentiation (**Fig. 4, 5**). This phenotype closely resembles that of mouse embryos lacking *Pofut1*, a core component of Notch signalling, in NC derivatives, where premature neuronal differentiation is accompanied by ENS progenitor depletion (Okamura and Saga, 2008). Although DAPT treatment appears to promote the generation of various enteric neuronal subtypes, including both cholinergic and nitrergic neurons (**Fig. 1, S3C**), it will be important to further determine whether Notch inhibition preferentially biases differentiation toward a specific neuronal differentiation trajectory, particularly given the reported expression of distinct Notch pathway components in later ENS populations (Morarach et al., 2021, Sander et al., 2003).

Similar observations have been reported in other neural progenitor systems (de la Pompa et al., 1997, Skeath and Carroll, 1992, Marklund et al., 2010, Trujillo-Paredes et al., 2016, Cheung et al., 2018), where Notch orchestrates differentiation timing to precursor pool size with the timely generation of derivatives. Interestingly, we found no evidence supporting the pro-gliogenic role of Notch signalling reported in the mouse ENS and peripheral nervous system (PNS) (Boesmans et al., 2021, Taylor et al., 2007, Mead and Yutzey, 2012, Okamura and Saga, 2008, Wakamatsu et al., 2000, Okubo et al., 2021, Wiszniak and Schwarz, 2019, Delfino-Machin et al., 2017). Instead, glial numbers appeared to increase following DAPT treatment, reflecting an accelerated overall rate of differentiation rather than a switch toward a gliogenic bias, as captured by our modelling, and suggesting species-specific differences or distinct temporal dynamics in human ENS development. It should be noted that our findings define the role of Notch signalling in the presence of GDNF, a key pro-ENS differentiation factor. Thus, investigating how Notch signalling interacts with extrinsic neurogenic cues to regulate human ENS development would be a key future research direction.

DAPT treatment also induced an earlier peak/potentiation in *DLL1/3/4* ligand expression, along with downregulation of *HES5/NOTCH1*, consistent with findings in *Notch1* and *RBP-Jk* mutant mice (de la Pompa et al., 1997). These results suggest that premature differentiation in our system may be mediated by a lateral inhibition feedback mechanism involving these Notch components, key fate determinants such as *SOX10* and *ASCL1*, and regulators of the cell cycle and other signalling pathways (e.g., Hedgehog, Wnt) (Okamura and Saga, 2008, Militi et al., 2023, Ngan et al., 2011). Given the dynamic temporal changes in the expression of Notch pathway components observed in the presence and absence of continuous DAPT treatment, a logical next step would be to define the stage-specific effects of Notch inhibition by targeting discrete windows of human ENS differentiation.

Our results further revealed that the accelerated generation of enteric neurons and glia in DAPT-treated cultures was accompanied by impaired migration of SOX10⁺ ENS progenitors (**Fig. 6**). This was reflected by reduced migration velocity and directional persistence, indicating that Notch inhibition directly influences the intrinsic migratory behaviour of ENS progenitors, consistent with previous studies implicating Notch signalling in cardiac, vagal, and trunk NC migration (Alhashem et al., 2022, Mead and Yutzey, 2012). An important avenue for future investigation will be to determine whether Notch signalling regulates collective ENS progenitor migration by coordinating cell–cell interactions and reciprocal proliferation status between leader and follower progenitors, as well as between ENS progenitors and neighbouring non-ENS cells, and how these interactions influence progenitor proliferation. Such a mechanism has been demonstrated in zebrafish trunk NC cells (Alhashem et al., 2022) and is further supported by single-cell transcriptomic analyses of wavefront ENS progenitors and their surrounding cells in the developing mouse embryo (Zhou et al., 2024).

Together, these findings support a model in which disruption of Notch signalling, either through direct mutation of pathway components (Ngan et al., 2011, Tang et al., 2017) or as a downstream consequence of attenuated ENS regulators (e.g. RET signalling (Vincent et al., 2023, Windster et al., 2025)), can lead to aganglionic phenotypes. This occurs via premature differentiation from the ENS progenitor state and impaired progenitor migration, two key features of abnormal ENS development in HSCR.

## METHODS AND MATERIALS

### hPSC culture and differentiation

Human iPSCs (WTC11 and Rainbow WTC11 (Miyaoka et al., 2014, El-Nachef et al., 2021) and H9 SOX10:GFP hESCs (Chambers et al., 2012) were routinely cultured in feeder-free conditions using Geltrex LDEV-Free reduced growth factor basement membrane matrix (A1413202, Thermo Fisher Scientific) and mTeSR medium (Stem Cell Technologies, #85850). Cells were passaged upon reaching approximately 80-90% confluency using PBS/EDTA (Invitrogen, #15575020) or ReleSR (Stem Cell Technologies, #05872).

Differentiation toward ENS cells was carried out as previously described (Gogolou et al., 2021). Briefly, hPSC cultures at 80-90% were dissociated into a single-cell suspension following treatment with Accutase (Merck Life Science, #A6964) and plated at a density of 30-50,000 cells/cm^2^ onto Geltrex-coated plates. Cells were cultured in DMEM/F12 (Merck, #D6421) supplemented with 1x N2 (Thermo Fisher, #17502048), 1x non-essential amino acids (Thermo Fisher, #11140050), 1x Glutamax (Thermo Fisher, #35050061), CHIR99021 (1μM; Tocris, #4423), SB431542 (2μM; Tocris, #1614/1), BMP4 (20ng/ml; Thermo Fisher, #PHC9533), and DMH-1 (1μM; Tocris, #4126/10) for 4 days. This was followed by the addition of all-trans retinoic acid (1μM; Merck, #R2625) for a further 2 days to generate vagal NC cells.

On D6, vagal NC cells were dissociated using Accutase and re-plated under non-adherent conditions to form free-floating spheres using Corning Ultra-Low attachment 6-well plates (Corning, #3471). Spheres were cultured for 3 days in medium containing 0.5x Neurobasal (Thermo Fisher, #21103049), 0.5x DMEM/F12 (Merck, #D6421), 1x non-essential amino acids (Thermo Fisher, #11140050), 1x Glutamax (Thermo Fisher, #35050061), 1x N2 (Thermo Fisher, #17502048), 1x B27 (Thermo Fisher, #17504044), CHIR99021 (3μM; Tocris, #4423), FGF2 (10 ng/ml; R&D Systems, #233-FB/CF), all-trans retinoic acid (1μM; Merck, #R2625), and Y-27632 2HCl (first 24 hrs; 10 μM; AdooQ Bioscience, #A11001).

Finally, D9 spheres were collected and cultured for up to 50 days on Geltrex-coated plates in ENS-inducing medium containing 1x BrainPhys (StemCell Tech, #05790), 1x non-essential amino acids (Thermo Fisher, #11140050), 1x Glutamax (Thermo Fisher, #35050061), 1x N2 (Thermo Fisher, #17502048), 1x B27 (Thermo Fisher, #17504044), Ascorbic Acid (200 μM; Sigma-Aldrich, #A8960), and GDNF (10 ng/ml; Peprotech, #450-10), in the presence of DAPT (10 μM dissolved in DMSO; Tocris, #A2634) or an equivalent volume of DMSO alone (0.1% v/v) as vehicle control, as described in the Results section. ENS medium was changed every other day (half medium change).

For plating, D9 spheres were collected into a 50ml conical tube and allowed to settle by gravity. Following careful removal of the medium, spheres from one well of the Ultra-Low attachment 6-well plate were resuspended and triturated in 4 ml of ENS induction medium, then plated at 1:4 and 1:8 dilutions in 12-well and 24-well plates, respectively.

For confocal microscopy, some wells of 12-well plates were fitted with 18 mm round glass coverslips (No. 1; VWR, #MENZCB00180RAC20). These were pre-treated overnight at 37°C with poly-L-ornithine solution (50 μg/ml in PBS, Thermo Fisher, A31804) prior to Geltrex-coating to facilitate adhesion.

To monitor migration of eGFP+ progenitors from plated spheres, eGFP-WTC-11 cells - sorted and cloned from the WTC-11 rainbow reporter cell line containing a modified Brainbow cassette (El-Nachef et al., 2021) - were differentiated into vagal NC cells. On D6, these cells were mixed at a 1:10 ratio with stage-matched, unlabelled cells derived from the parental WTC-11 line and subjected to sphere formation for 3 days. Spheres were then re-plated under ENS-inducing conditions and live-imaged for 75 hours.

### Quantitative real-time PCR

Total RNA was extracted using the Norgen total RNA purification kit (Norgen BioTek, #48200), following the manufacturer’s instructions. Samples from undifferentiated hPSCs were collected as cell pellets immediately prior to initiating differentiation, representing the starting population used to generate the ENS cultures. RNA was extracted from these samples and analysed alongside the differentiated samples within the same qPCR experiment. cDNA synthesis was performed using the High-Capacity cDNA Reverse Transcription Kit (Thermo Fisher, #4368813), according to the manufacturer’s instructions. Quantitative real-time PCR was carried out using the QuantStudio 12 K Flex (Applied Biosystems) thermocycler in combination with the Roche Universal Probe Library system and TaqMan™ Fast Universal PCR Master Mix no AmpErase™ UNG (Applied Biosystems, #4366072). *GAPDH* was used as the housekeeping gene, serving as both a positive amplification control and reference gene for relative quantification normalization. Primer sequences (UPL probe number) were as follows: *GAPDH*: 5’-agccacatcgctcagacac-3’ and 5’-gcccaatacgaccaaatcc-3’ (#60); *SOX10*: 5’-ggctcccccatgtcagat-3’ and 5’- ctgtcttcggggtggttg-3’ (#21); *S100B*: 5’-gagctttcccatttcttagagga-3’ and 5’- gaagtcacattcgccgtctc-3’ (#47); *ASCL1*: 5’-cgacttcaccaactggttctg-3 and’ 5’- atgcaggttgtgcgatca-3’ (#38); *PHOX2B*: 5’-ctaccccgacatctacactcg-3’ and 5’-ctcctgcttgcgaaacttg-3’ (#17); *PRPH*: 5’-aagacgactgtgcctgaggt-3’ and 5’- tgctccttctgggactctgt-3’ (#10); *RET*: 5’-catcaggggtagcgaggtt-3’ and 5’- gggaaaggatgtgaaaaca-3’ (#17); *TRKC*: 5’- ccgtacgagagggtgacaat-3’ and 5’- tggtccagttcagattggtct-3’ (#21); *SST*: 5’-accccagactccgtcagttt-3’ and 5’-acagcagctctgccaagaag-3’ (#38); *CHAT:* 5’-cagccctgatgccttcat-3’ and 5’-cagtcttcgatggagcctgt-3’ (#78); *TH*: 5’-acgccaaggacaagctca-3’ and 5’- agcgtgtacgggtcgaact-3’ (#42); *HTR2A*: 5’- tgatgtcacttgccatagctg-3’ and 5’-caggtaaatccagactgcacaa-3’ (#3). For Notch signalling-associated gene expression analysis, the following predesigned primer/probe mixtures (PrimeTime™ Gene Expression Assays, IDT) were used (assay IDs): *GAPDH*: Hs.PT.39a.22214836; *NOTCH1:* Hs.PT.58.23074795; *NOTCH2:* Hs.PT.58.28296019; *HES1:* Hs.PT.58.4181121; *HES5:* Hs.PT.58.14966721; *JAG1*: Hs.PT.56a.4972610; *JAG2*: Hs.PT.58.3493191; *DLL1*: Hs.PT.58.41063402; *DLL3*: Hs.PT.58.21516515; *DLL4*: Hs.PT.58.3416363. Graphs were generated using GraphPad Prism (GraphPad Software), which was also used for statistical analysis.

### Immunofluorescence / Image analysis

Cells were fixed in 4% paraformaldehyde (J61899.AP, VWR) for 10 minutes at room temperature. They were then permeabilised for 15 minutes at room temperature using 0.1-0.5% Triton X-100 in PBS (PBST), followed by incubation (1 hour at room temperature or overnight at 4°C) in blocking buffer comprising 3% donkey serum (Jackson ImmunoResearch, #017-000-121) / 1% BSA in PBST. Primary and secondary antibodies, diluted in blocking buffer at the appropriate working concentrations (see below), were then added for 24-48 hours at 4°C with gentle shaking on a rocking platform. Following primary antibody incubation, samples were washed at least three times for 5 minutes each with PBS, after which fluorophore-conjugated secondary antibody solution was added. Cells were incubated for 1-2 hours at room temperature or overnight at 4°C in the dark. After secondary antibody incubation and nuclear counterstaining using Hoechst 33342 (Thermo Fisher Scientific, #H3570), cells were washed three times with PBS prior to analysis. Fixed/stained cells on coverslips were mounted on 75×25 mm glass slides (VWR, Menzel Gläser, SuperFrost Plus) using VECTASHIELD mounting medium (2B Scientific, #H-1700-10) and left to dry in the dark before imaging or storage at 4°C.

Fluorescent images were acquired using either the InCell Analyser 2200 system (GE Healthcare) or a LSM880 confocal system with Airyscan detectors (Zeiss). Images were first processed using ZEN (ZEISS) software, in the case of confocal imaging, and then analysed in Fiji (Schindelin et al., 2012). Identical brightness and contrast settings were applied across all images to enable comparison between different treatments. Samples incubated with secondary antibody solution only were used throughout to establish a threshold for positive signal, thereby minimising labelling artefacts introduced by fluorophore-conjugated secondary antibodies.

The following primary antibodies and dilutions were used: Anti-SOX10 (1:500; CST #89356); anti-SOX10 (1:200; R&D systems #AF2864); anti-PERIPHERIN (1:60-1:100; Millipore Sigma #AB1530); anti-PERIPHERIN (1:200; Novus Biolog. #NBP1-05423); anti-ASCL1 (1:100; Abcam #ab211327); anti-PGP9.5 (1:1000; Abcam #ab108986); anti-ChAT (1:200; Millipore Sigma #AB144P); anti-HuC/HuD (1:60-1:100; Invitrogen #A-21271); anti-PHOX2B (1:50; Santa Cruz #sc-376997); anti-S100 (β-Subunit) (1:1000; Millipore Sigma #S2532); anti-S100 (1:50-1:250; Dako Omnis #Z0311); anti-beta III Tubulin(TuJ1) (1:1000; Abcam #ab78078); anti-beta III Tubulin(TuJ1) (1:1000; BioLegend #PRB-435P);anti-TRKC (1:500-1000; CST #3376); anti-Substance P (1:100; R&D Systems #MAB4375); anti-CD49d (1:1000; BioLegend #304302); anti-nNOS (1:400; Invitrogen #61-7000).

Protein expression was quantified using either: (1) CellProfiler (Carpenter et al., 2006) with custom pipelines for nuclear segmentation and intensity measurements; or (2) CellPose (Stringer et al., 2021) for nuclear segmentation using a custom-trained model, refined through iterative human-in-the-loop training, with segmentation masks imported into PickCells (https://pickcellslab.frama.io/docs/) for fluorescence intensity analysis. The latter approach was used to quantify HuC/D/SOX10/S100β-positive cells in ENS cultures.

Nearest neighbour and internuclear edge distance analyses were performed using PickCells, as described previously (Punovuori et al., 2019). Cells were segmented using CellPose based on Hoechst 33342 nuclear staining, and segmentation masks were imported into PickCells. Nearest neighbours for each HUC/D^+^ cell were identified using Delaunay triangulation. Given the limited cytoplasmic area, nuclei were used as proxies for cell positions. Distance between neighbouring cell membranes were calculated for each HUC/D^+^ nucleus within a radius of 50 μm.

Scratch assay and wound closure analyses were performed on time-lapse data (see section below) using a previously published ImageJ plugin (Suarez-Arnedo et al., 2020). The wound region at each time point was defined by a segmentation algorithm, and cell migration was quantified by measuring leading-edge progression over time. Percentage wound closure was calculated for each time point and condition relative to control (DMSO, set to 1). Scratch area was measured in μm^2^.

Statistical significance was calculated using GraphPad Prism (GraphPad Software).

### Live imaging

Time-lapse microscopy was performed at 37°C and 5% CO_2_ using either a Nikon Biostation CT or a Nikon Ti inverted microscope equipped with a stage-top incubation chamber and a DS-Fi3 colour camera. To monitor migration of eGFP+ progenitors from plated spheres, the Nikon Biostation CT was used, with images captured every 10 minutes for 75 h using a 10x air objective. Images were acquired in both phase-contrast and green fluorescence channels. Image stacks were compiled in CL Quant (Nikon) and exported to Fiji (Schindelin et al., 2012) for analysis.

For scratch assay analysis, wound closure was imaged using either a Nikon Biostation CT or a Nikon Ti inverted microscope every 10 minutes for 24 hours using a 10 x air objective (either Plan Fluor Ph1 for phase contrast or Plan Apo λ for brightfield). Imaging positions were selected manually and 3-5 positions across 1-3 wells per condition and per experiment were monitored. Images were initially processed using Nikon software; where necessary, multiple fields of view were stitched to generate images spanning the full wound width, then exported to Fiji for analysis.

### Scratch assay

D9 spheres were plated into Geltrex-coated 24-well plates in ENS induction medium (DMSO control or 10μM DAPT or GDNF and Ascorbic Acid) and incubated at 37°C with 5% CO_2_ until a confluent monolayer formed. On D14 of differentiation, a scratch assay was performed by manually removing cells from the centre of each well using a plastic 200 μl pipette tip. Wells were then washed three times with 1x PBS to remove detached cells, and fresh ENS medium containing either DMSO or 10μM DAPT was added. Plates were incubated at 37°C and 5% CO_2_ in either the Biostation CT or a stage-top incubation chamber on a Nikon Ti inverted microscope for 24 hours, with images acquired every 10 minutes.

### Quantification of cell migration dynamics

Single-cell migration dynamics were quantified using the Manual Tracking plugin in Fiji. Cell trajectories were analysed over a 6-hour observation period starting from 3 hours post-wounding. For each condition (DMSO/DAPT), 10 individual cells along the wound edge were manually traced per field of view (3 fields of view per condition). The experiment was conducted across 2 independent biological replicates, yielding a total of n=60 tracked cells per experimental group (30 cells per biological replicate). Spatial scale was maintained in pixel coordinates (1 pixel unit). To characterise cell movement, three key trajectory parameters were calculated from x, y coordinates across time: migration velocity, Y-axis Displacement and the straightness index (SI). Migration Velocity, defined as the mean movement speed of each individual cell, was calculated over the entire tracking duration and subsequently averaged per field of view. Y-axis displacement, defined as the net spatial displacement along the y-axis (perpendicular to the initial wound edge), was measured to evaluate directional progress into the wound area. SI was quantified as the ratio of net direct displacement (D) to the total trajectory path length (L)

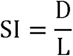

An SI value approaching 1 indicates linear, persistent migration, whereas values approaching 0 reflect random movement.

### Mathematical modelling

We modelled the dynamics of ENS progenitor differentiation following the dissociation and replating of D9 neural spheroids and further culture for an additional 13 days, until D22 of differentiation, with (DAPT-treated) or without (DMSO control) NOTCH-inhibition (NOTCHi) (**Fig. 5C**). Our mean-field model comprises a set of coupled ordinary differential equations (ODEs), which reflect our assumptions regarding cell-state transitions, proliferation, and death (**Fig. 5A, B**). These equations govern the evolution in time (*t*) of the numbers of SOX10^+^/S100β^−^/HUC/D^−^ progenitors (*P*), SOX10^−^/S100β^−^/HUC/D^+^ neurons (*N*), SOX10^+^/S100β^+^/HUC/D^−^ glia (*G*), and SOX10^−^/S100β^−^/HUC/D^−^ unidentified cells (*U*).

We assumed that ENS progenitors differentiate irreversibly with per-capita rate *k*(*t*) (**Fig. 5A**), giving rise to neurons and glia; the neuronal and glial shares of differentiation are set by a bias *ρ*(*t*) and its complement 1- *ρ*(*t*), respectively. We assumed that ENS progenitors proliferate with constant net per-capita rate *α*, which captures the net effect of division and death on cell number (**Fig. 5B**); that glia, given their low reported rate of division (Joseph et al., 2011, Laddach et al., 2023), do not appreciably self-renew; and that neurons are quiescent (Morarach et al., 2021). The SOX10^−^/S100β^−^/HUC/D^−^ unidentified cells were treated as a co-emerging population that is not derived from the ENS progenitors but is present in the cultures, and evolves with its own net per-capita rate *α_U_*. We assume the cells within the unidentified population represent mesenchymal-like cell types derived from day 6 vagal NC and/or cranial/cardiac NC contaminants.

Accounting for these processes led to the following ODE system describing cell population dynamics during differentiation with or without Notch inhibition:

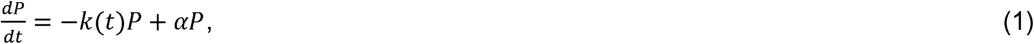

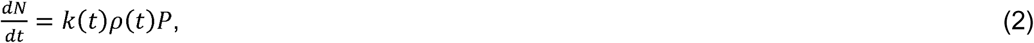

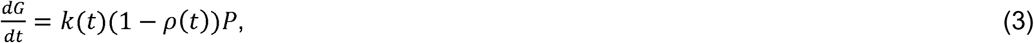

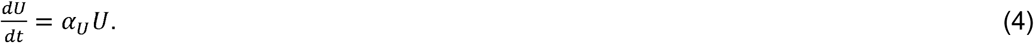

We assumed that immediately following replating at D9, no neurons or glia were present, so that *N*(9) = 0 and *G*(9) = 0. Furthermore, we assumed that an undetermined proportion *P*_0_ of cells at this time point were ENS progenitors, with the remainder unidentified. Finally, we scaled all dependent variables by the total cell number at D9, so that *P*(9) = *P*_0_ and *U*(9) = 1 – *P*_0_. We assumed the same model structure with or without Notch inhibition.

We tested four distinct scenarios: constant differentiation rate and lineage biases (CC); constant rate and time-varying biases (CV); time-varying rate and constant biases (VC); and time-varying rate and biases (VV). The functional forms of □ and *ρ* are given below:

**Constant:**

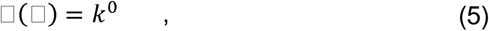

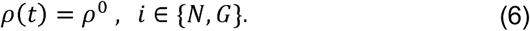

**Linear:**

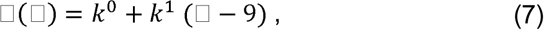

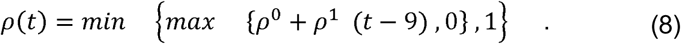

The min/max expression in (8) ensures that the neuronal lineage bias does not fall below 0 or exceed 1.

Equations (1)-(8) were solved numerically in Python; see the ‘Data and resource availability’ section for details on how to download our code.

### Parameter inference and model selection

As most of our model parameters cannot be measured directly, they must be inferred from the data. Because our model predicts cell-type proportions rather than absolute numbers, and because the field-of-view counts are substantially over-dispersed relative to binomial sampling (varying between biological replicates by considerably more than sampling alone would predict), we inferred parameters using a compositional likelihood evaluated directly on the observed counts, conditioning on the total number of cells counted in each field.

For each timepoint *j* and biological replicate *r*, let *π_P_*(*t_j_*), *π_N_*(*t_j_*), *π_G_*(*t_j_*), and *π_U_*(*t_j_*) denote the model-predicted proportions of progenitors, neurons, glia and unidentified cells, obtained by normalising the solution of equations (1)–(4) by the total cell number. In the neuronal field, the number of HUC/D⁺ cells was modelled with a beta-binomial distribution,

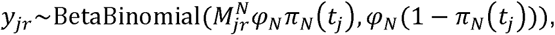

where 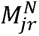 is the total number of cells scored in that field and *y_jr_* the number that are HUC/D⁺. In the SOX10/S100β field, the joint counts of SOX10⁺S100β⁻ progenitors (*n_P_*), S100β⁺ glia (*n_G_*) and the remaining cells 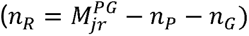 were modelled with a Dirichlet–multinomial distribution,

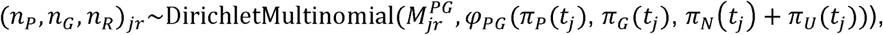

where 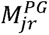 is the total number of cells scored in that field and the remainder category corresponds to neurons and unidentified cells, which are not separately marked in this field. Here *φ_N_* and *φ_PG_* are field-level concentration (precision) parameters that capture the between-replicate over-dispersion: larger values approach ordinary binomial or multinomial sampling, whereas smaller values admit greater replicate-to-replicate variability about the predicted proportions. Both were inferred jointly with the dynamical parameters. Since each distribution conditions on the observed field total, this likelihood compares predicted and observed *proportions* directly and does not depend on the unknown initial cell number.

We sampled posterior distributions using the No-U-Turn sampler (Hoffman and Gelman, 2014), a self-tuning variant of Hamiltonian Monte Carlo implemented in NumPyro, running four independent chains per scenario. Weakly-informative priors were placed on the time-varying rate and bias gradients to regularise the control condition, in which these gradients are only weakly identified. Convergence was assessed using the rank-normalised R-hat statistic and bulk and tail effective sample sizes (Vehtari, 2021); across all fitted parameters R-hat did not exceed 1.013, and we confirmed that an independent affine-invariant ensemble sampler (emcee;(Foreman-Mackey et al., 2013)) recovered indistinguishable posterior medians. The four scenarios were compared by leave-one-biological-replicate-out cross-validation, quantified by the expected log predictive density (Vehtari et al., 2017), which evaluates each model’s ability to predict held-out experimental replicates rather than its in-sample fit. Parameter inference and model selection were performed using the experimental datasets shown in **Fig. 4E** and **Fig. 4G**.

We set parameter priors as follows:

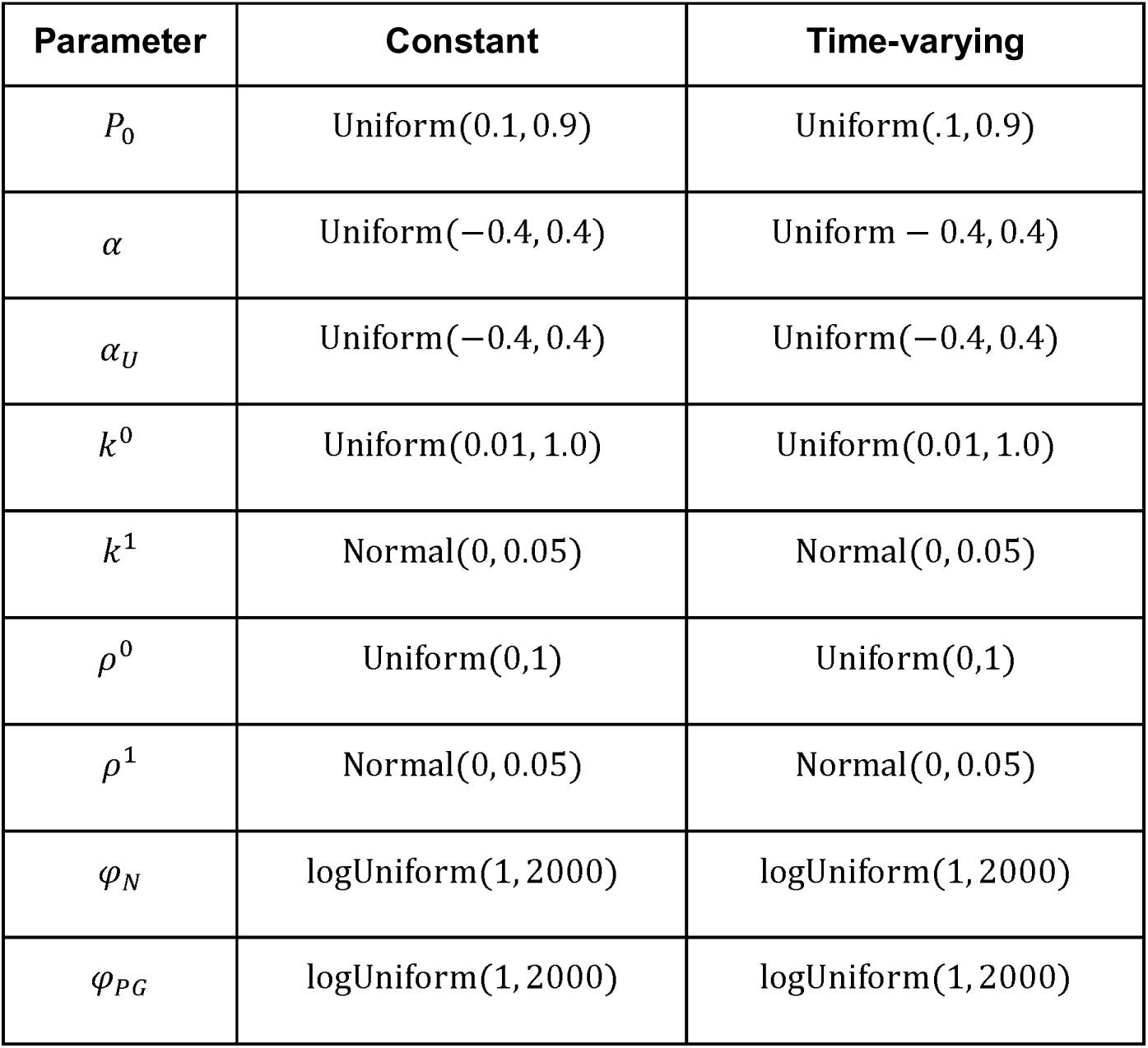

The parameter *P_0_* was only inferred for –NOTCHi differentiation, this was then used for +NOTCH inference. A summary of inferred parameters can be found in **Supplemental Table S1**.

### Comparison of posterior parameter distributions

Between-condition differences in the inferred parameters were characterised by comparing their marginal posterior distributions directly, using the posterior medians and 90% credible intervals summarised in **Supplemental Table S1** and shown in **Fig. 5E–H, K–N**. We regarded a parameter as differing between the control and Notch-inhibited conditions when its marginal posteriors showed limited overlap, for example non-overlapping 90% credible intervals, and as essentially unchanged when the intervals overlapped substantially. We did not apply a formal significance test to the posterior samples, because with a large number of posterior draws such tests primarily reflect the precision with which each posterior is estimated rather than the magnitude of any biological difference.

### Data and resource availability

All code used for simulations, model fitting, and statistical analysis, together with datasets used for modelling, are publicly available on GitHub (https://github.com/AlexFletcher/gogolou_2026_notch).

## Acknowledgements

We thank Darren Robinson and Nick Van Hateren for assistance with imaging, which was performed at the Wolfson Light Microscopy Facility, University of Sheffield. We also thank Tom Frith, Chris Price and Rebecca Lea for technical support with the live imaging experiments. Finally, we are grateful to Billy Douglas for his continuous support and Peter Andrews, Conor McCann and Celin Souilhol for critical reading of the manuscript.

## Competing interests

The authors declare no competing or financial interests.

## Funding

This research was funded in part by the UKRI Medical Research Council (MR/V002163/1 and MR/Y013476/1; AT, PWA), the European Union Horizon 2020 Framework Programme (H2020-EU.1.2.2; project 824070; AT, PWA, AG), the UKRI Biotechnology and Biological Sciences Research Council (White Rose Mechanistic Biology DTP studentship; NS], the Wellcome Trust (Sir Henry Wellcome Postdoctoral Fellowship 224070/Z/21/Z; SES), and the University of Sheffield (Strategic Research Fellowship in the Physics of Life and Quantitative Biology; SES).

## Author contributions

**Conceptualization:** AG, NS, SES, AGF, AT

**Methodology:** AG, NS, SES, AGF

**Software:** GB, NS

**Validation:** AG, NS, SES

**Formal analysis:** AG, NS, SES

**Investigation:** AG, NS, SES

**Data Curation:** AG, NS, SES

**Writing – original draft preparation:** AG, NS, SES, AGF, AT

**Writing – review and editing:** AG, NS, GB, SES, AGF, AT

**Visualization:** AG, NS, SES

**Supervision:** SES, AGF, AT

**Project administration:** AT

**Funding acquisition:** SES, AGF, AT

---

**Figure S1. Emergence of human enteric neurons *in vitro*.** (A) Schematic of the experimental strategy used to monitor ENS differentiation following sphere plating. GFP-expressing (Rainbow WTC-11) and unlabelled (WTC11) cells were differentiated to vagal NC and mixed at a 1:10 ratio prior to sphere formation. On day 9, spheres were plated under ENS-inducing conditions and live-imaged to track migration of GFP^+^ progenitor cells. (B) Representative snapshots from time-lapse microscopy (brightfield and fluorescence) showing neurons migrating out of spheres at the indicated hours (h) after plating on day 9 (D9). Insets show magnified areas. Images were obtained every 10 minutes for 75 hours. (C) Immunofluorescence analysis of ASCL1, PHOX2B and PERIPHERIN in day 50 ENS cultures. The arrow indicates a triple-positive cell. (D-F) Immunofluorescence analysis of the indicated enteric neuronal markers in day 50 ENS cultures. Arrows indicate double-positive cells. Scale bars: 30 μm.

**Figure S2. In vitro differentiation of SOX10:GFP hESCs toward ENS cell types.** Immunofluorescence analysis of hESC-derived ENS cultures generated using the H9SOX10:GFP reporter cell line. (A-C) Expression of PERIPHERIN, HUC/D (A) TUJ1, SOX10:GFP (B) and NF-H, SOX10:GFP and S100β (C) in day 22 cultures. (D, E) Expression of PERIPHERIN, HUC/D, SOX10:GFP (D) and RET, SOX10:GFP (E) in day 38 cultures. Insets show magnified regions. (F-I) Expression of TUJ1 and the neural subtype marker CALRETININ (F), PERIPHERIN (G), TH (H) and ChAT (I) in day 55 ENS cultures. Scale bars=50 μm. D, days.

**Figure S3. Impact of Notch inhibition on ENS progenitor differentiation.** qPCR analysis of *HES1* (A) and *HES5* (B) expression in day 22 ENS cultures derived from H9SOX10:GFP hESCs and treated with increasing concentrations of DAPT (10, 20, and 50 μM) or a vehicle control (DMSO, matched to the 50 μM DAPT treatment). Relative gene expression was normalised to the vehicle control (set to 1). Data are presented as mean ± s.d. from three biological replicates (n = 3). (C) qPCR analysis of indicated ENS markers in day 22 cultures. Data are shown as mean values ± s.e.m (n=5). Different symbols indicate independent experiments. Statistical analysis: two-way ANOVA followed by Sidak’s post-hoc multiple comparisons test; *P <0.05, **P <0.01, ***P<0.001. Only statistically significant changes are indicated.

**Figure S4. Posterior predictive checks and parameter distributions for ENS progenitor differentiation in DMSO control conditions.** (A-D) Numerical simulations (lines) of progenitor (purple), neuronal (yellow), and glial (blue) fractions for models where differentiation rate is constant (A, C) or time-varying (B, D) and lineage bias is constant (A, B) or time-varying (C, D), respectively, using 50 samples drawn from the posterior distribution inferred from empirical data on ENS progenitor differentiation dynamics (markers and boxplots; see also **Fig 4E, G,** DMSO). Open circles indicate biological replicates, box plots show the median and interquartile range of the technical replicate data, and lines denote simulations. (E) Posterior marginal distributions of inferred parameters for the initial progenitor fraction *P_0_*, the net per-capita growth rate of ENS progenitors α, the net per-capita rate of the unidentified population α*_U_*, the initial differentiation rate *k*^0^, the differentiation rate gradient k^1^, the initial lineage bias *ρ*, and the neuronal lineage bias gradient, *ρ*^1^.

**Figure S5. Posterior predictive checks and parameter distributions for ENS progenitor differentiation in DAPT treatment conditions.** (A-D) Numerical simulations (lines) of progenitor (purple), neuronal (yellow), and glial (blue) fractions for models where differentiation rate is constant (A, C) or time-varying (B, D) and lineage bias is constant (A, B) or time-varying (C, D), respectively, using 50 samples drawn from the posterior distribution inferred from empirical data on ENS progenitor differentiation dynamics under NOTCH inhibition (markers and boxplots; see also **Fig 4E, G,** DAPT). Open circles indicate biological replicates, box plots show the median and interquartile range of the technical replicate data, and lines denote simulations. (E) Posterior marginal distributions of inferred parameters: the net per-capita growth rate of ENS progenitors α, the net per-capita rate of the unidentified population *α_U_*, the initial differentiation rate *k*^0^, the differentiation rate gradient *k*^1^, the initial lineage bias *ρ*, and the neuronal lineage bias gradient *ρ*^1^.

**Figure S6. Cell type proportions normalised to the ENS compartment under control and Notch-inhibited conditions.** Proportions of ENS cell types expressed relative to the ENS compartment alone (progenitors + neurons + glia), rather than to all cells scored, for DMSO control (top row) and DAPT treatment (bottom row). Leftmost panels show the ENS compartment as a fraction of all cells; remaining panels show progenitor (purple), neuronal (yellow) and glial (blue) proportions within that compartment. Open circles indicate biological replicates, obtained by combining the HuC/D and SOX10/S100β immunofluorescence fields for each replicate; horizontal bars denote their means. Lines denote simulations using median posterior parameter values for models where differentiation rate is constant (black) or time-varying (red) and lineage bias is constant (solid) or time-varying (dashed); see also Fig. 4E, G. Because the unidentified population is dynamically uncoupled from the ENS lineages, these normalised proportions are independent of *P*_0_ and *α_U_*, and are obtained from the same posterior distributions as Fig. 5 without refitting.

**Figure S7. Effect of Notch inhibition on human ENS progenitor migration.** (A) Percentage of closure area. Data are presented as mean ± s.e.m, n=3 independent experiments with 3-5 fields scored per condition. Fields were sampled from 1-3 wells per experiment. (B) Scratch area in μm^2^. Data are presented as mean ± s.e.m, n=3 independent experiments with 3-5 fields scored per condition. Fields were sampled from 1-3 wells per experiment. (C) Representative brightfield time-lapse images of DMSO and DAPT treated ENS cultures captured at 3, 6 and 9 hour hours post-wounding. Coloured lines/dot show individual tracked cell trajectories.

## Notes

### Competing Interest Statement

The authors have declared no competing interest.

### Summary of Updates

The revised version includes new data and text changes aiming to address reviewer comments.

