## Supplementary figures and images for "Notch signalling governs human enteric nervous system progenitor dynamics" (from another version of this preprint: version 1, 2026-05-04)

### Fig S1

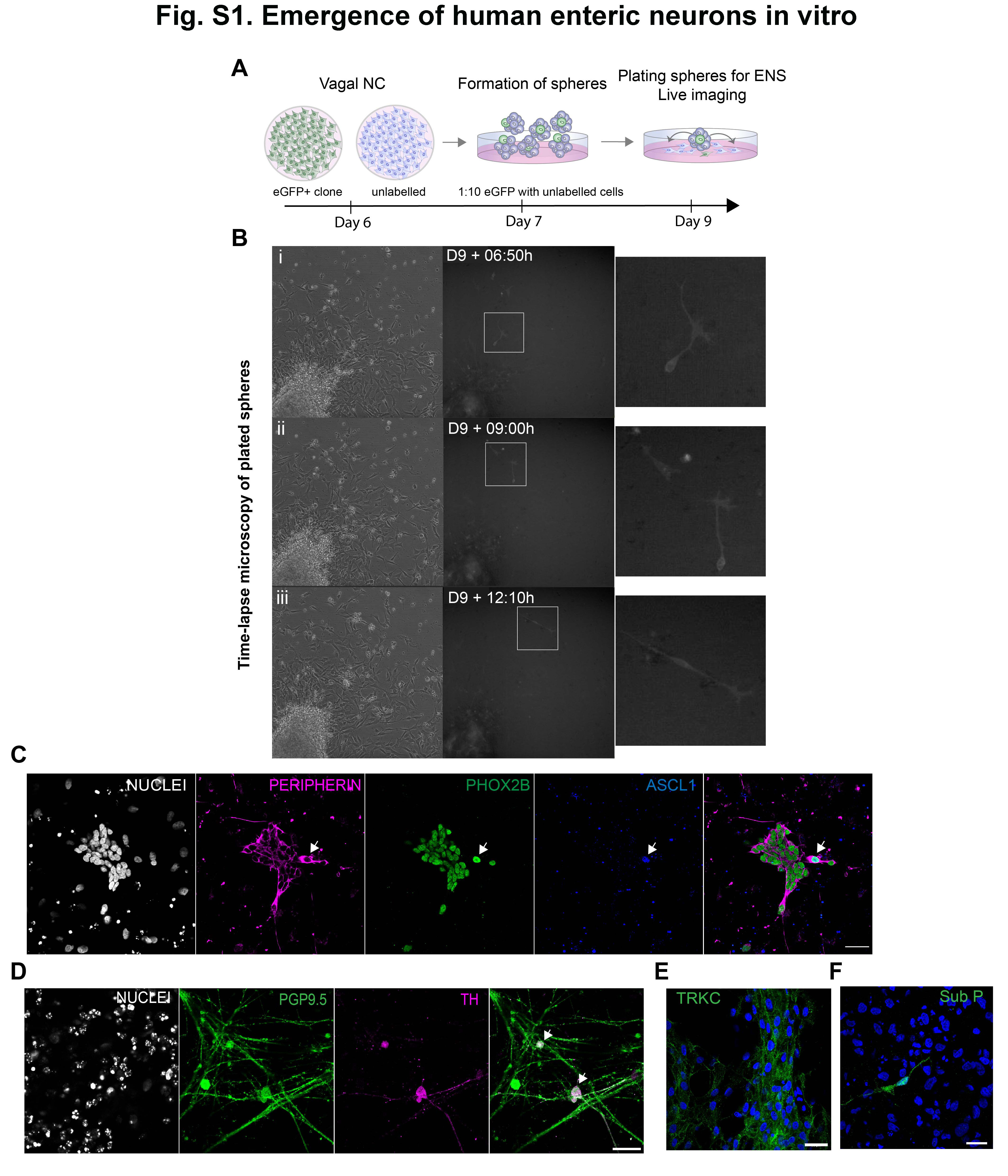

### Fig S2

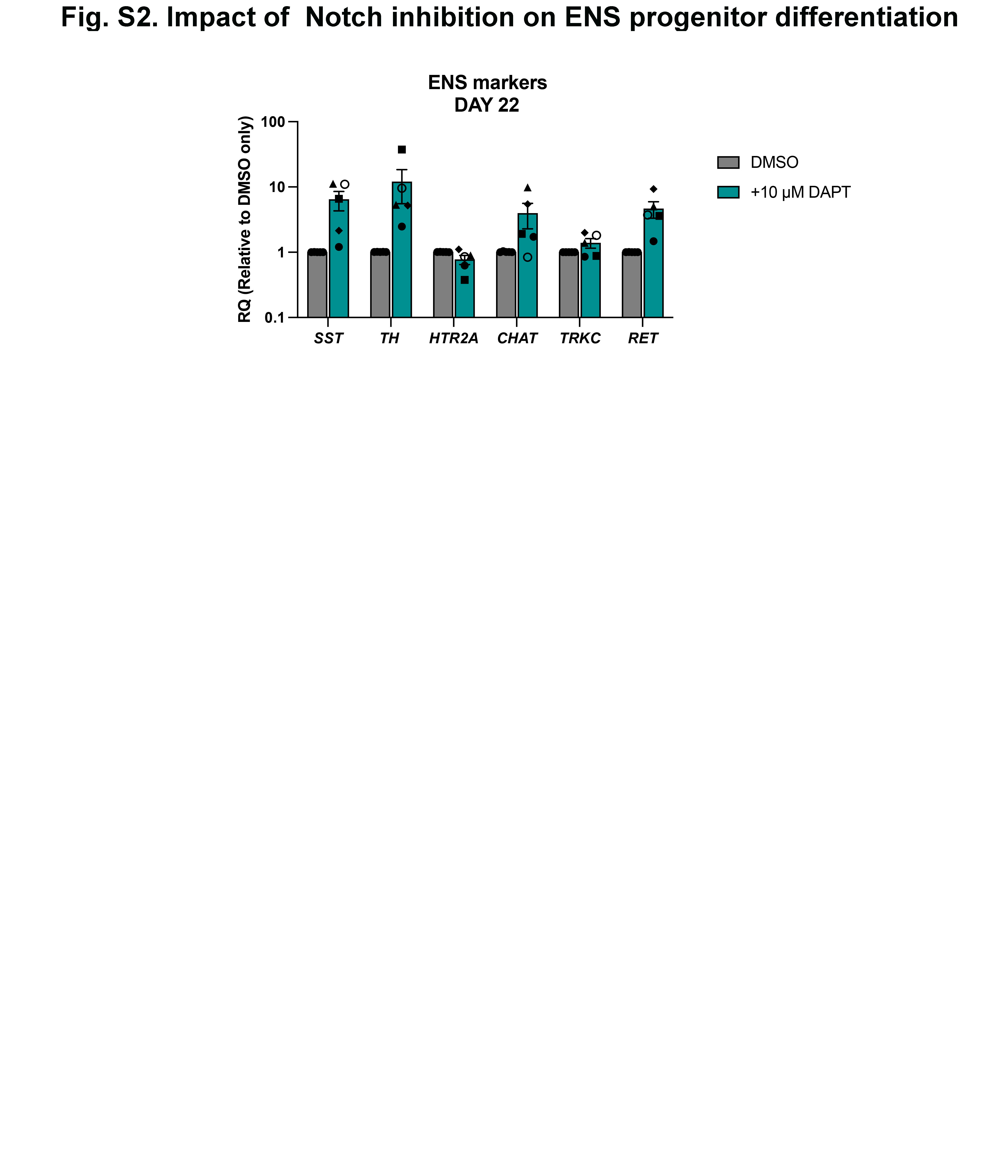

### Fig S3

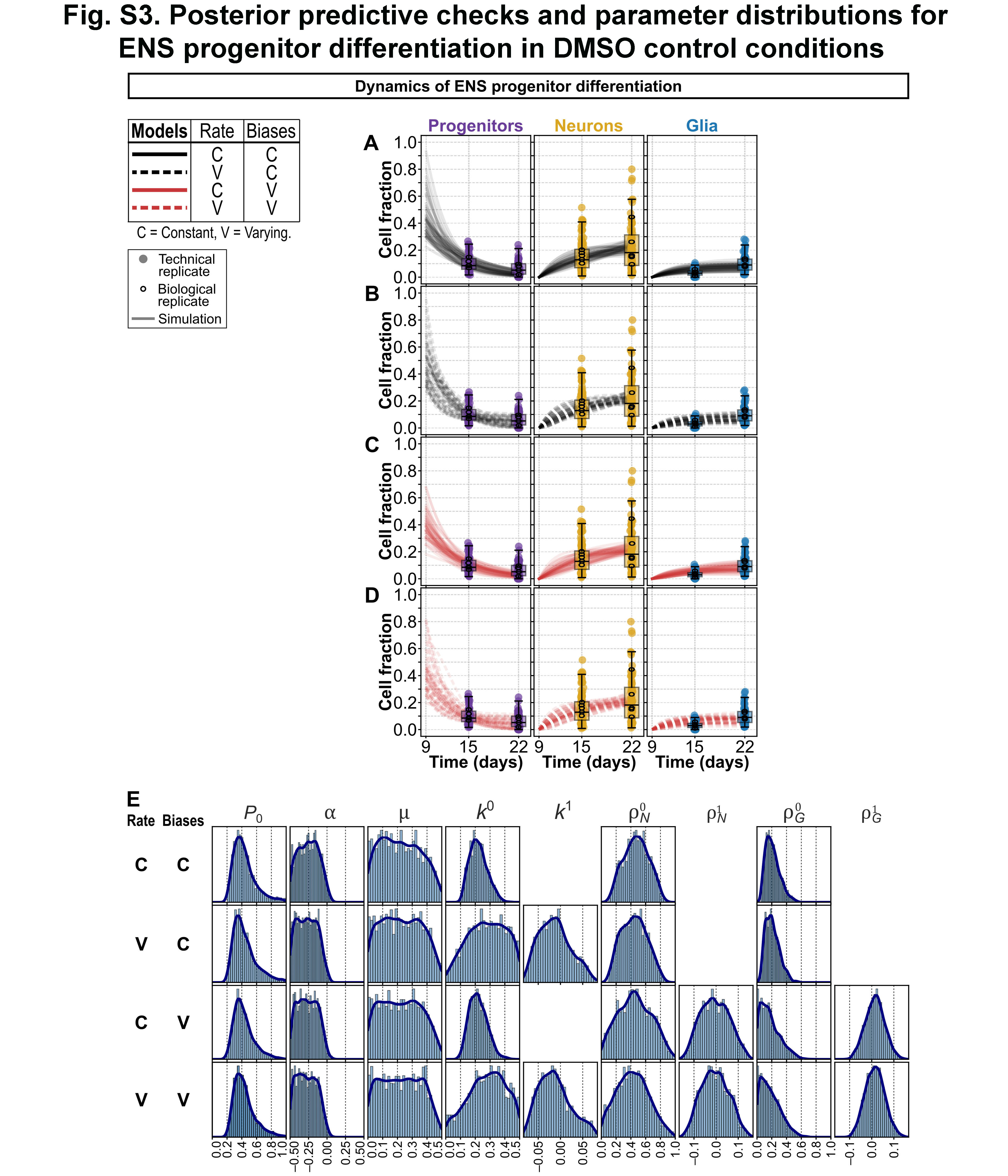

### Fig S4

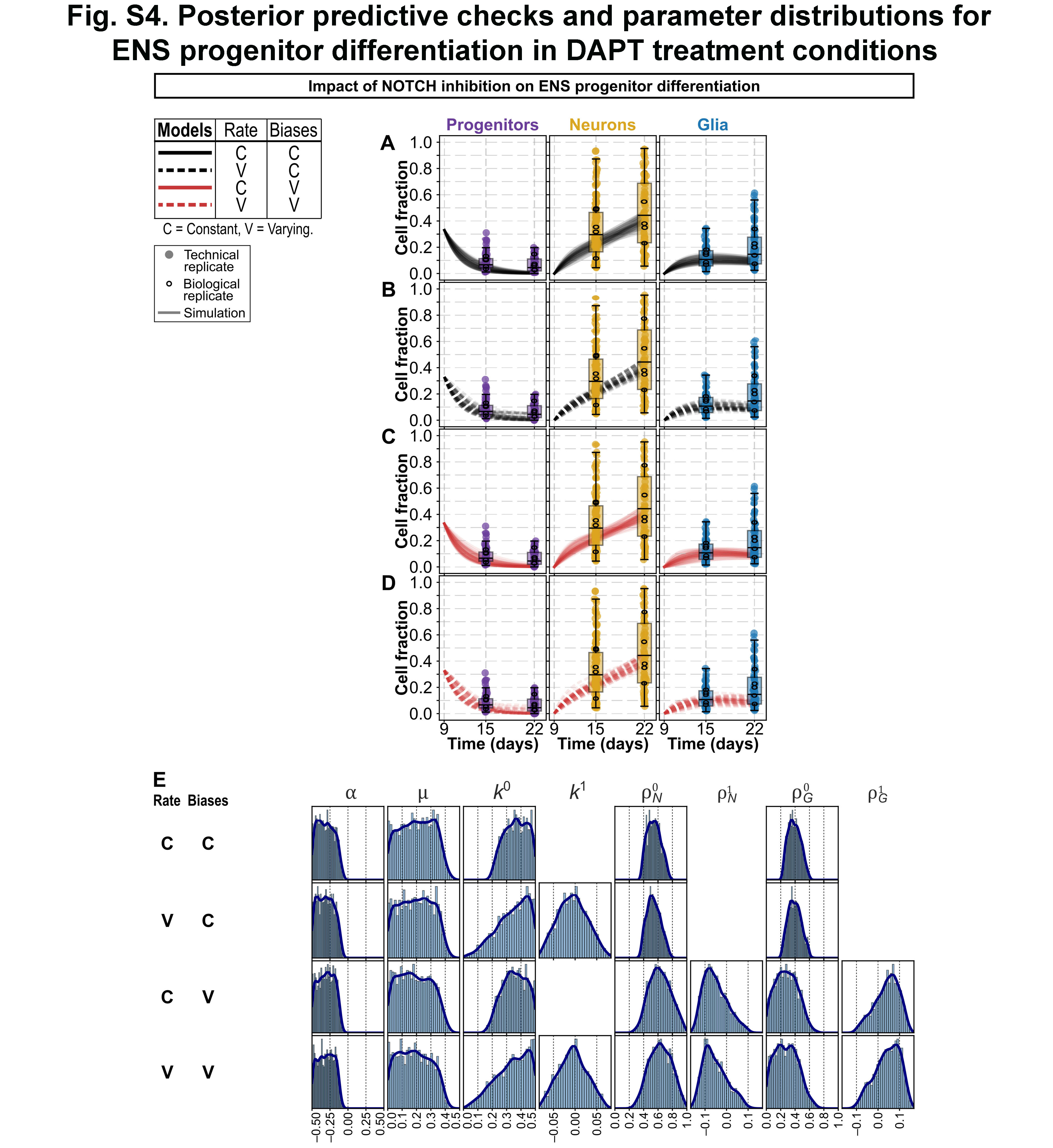

### Fig S5

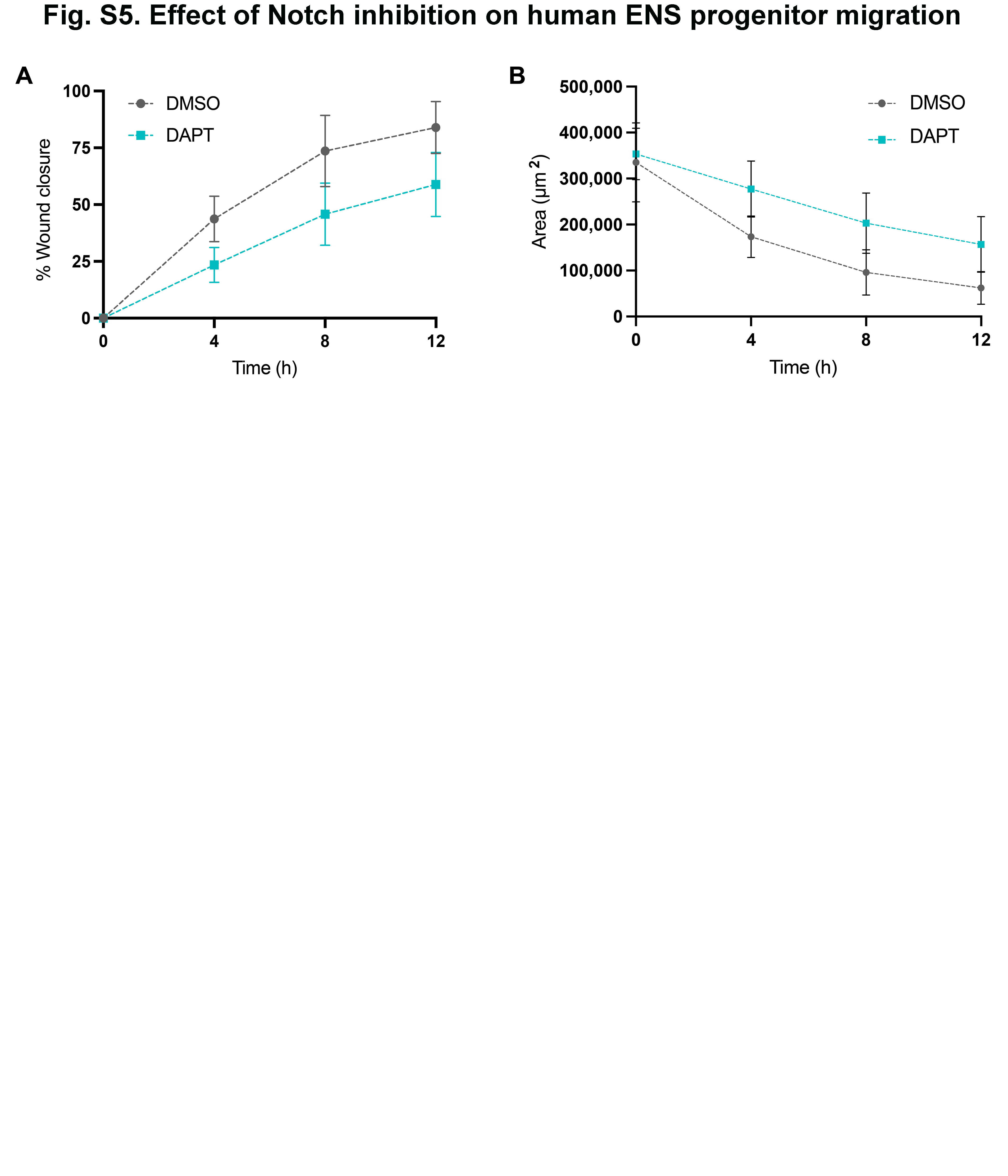
